# The splicing isoform Foxp3Δ2 releases the autoinhibitory conformation and differentially regulates tTregs and pTregs homeostasis

**DOI:** 10.1101/2023.01.09.523191

**Authors:** Qianchong Gu, Xiufeng Zhao, Jie Guo, Wei Xu, Jianhua Zhang, Wei Zhang, Fuping Zhang, Baidong Hou, Xuyu Zhou

**Affiliations:** CAS Key Laboratory of Pathogenic Microbiology and Immunology, Institute of Microbiology, Chinese Academy of Science (CAS), Beijing 100101, China; Department of Savaid Medical School, University of Chinese Academy of Sciences, Beijing 100049, China; Key Laboratory of Infection and Immunity, Institute of Biophysics, Chinese Academy of Sciences(CAS), Beijing 100101, China

**Keywords:** Foxp3, alternative splicing, conformation regulation

## Abstract

Foxp3 is the master transcription factor for the development and function of regulatory T cells (Tregs). So far, little is known about whether the conformation change in Foxp3 could impact the Tregs biology. Alternative splicing of human Foxp3 results in the expression of two major isoforms: the full-length protein or an exon 2-deleted protein (Foxp3Δ2). Here, AlphaFold2 structure predictions and *in vitro* experiments demonstrated that the N-terminal domain of Foxp3 inhibits DNA binding by moving toward the C-terminus and that this movement is mediated by exon 2. Consequently, we generated exon 2 deficient mice and found Foxp3Δ2-bearing Tregs in the peripheral lymphoid organ were less sensitive to TCR due to the enhanced binding of Foxp3Δ2 to the *Batf* promoter and were unsusceptible to IL-2. In contrast, among RORγt^+^ Tregs in the large intestine, Foxp3Δ2 Tregs expressed much more RORγt-related genes, and more strikingly, the deletion of exon 2 of Foxp3 conferred a competitive advantage over WT RORγt^+^ Tregs. Together, our results reveal that alternative splicing of exon 2 generates a constitutively active form of Foxp3, which plays a differential role in regulating tTregs and pTregs homeostasis.

**Highlights:**

1. Foxp3Δ2 broke the inhibitory loop and generated constitutive DNA-binding activity.
2. Foxp3 isoforms differentially regulate tTregs and pTregs homeostasis
3. Foxp3Δ2-bearing Tregs in the peripheral lymphoid organ were less sensitive to TCR and were unsusceptible to IL-2
4. Foxp3Δ2 RORγt^+^ pTregs benefited them for better adapting to the gut environmental conditions

## Introduction

Regulatory T cells (Tregs), a small subset of CD4 T cells, are essential in maintaining immune self-tolerance through dominant suppression mechanisms. Numerous preclinical studies have demonstrated that various autoimmune diseases are associated with the defection of Tregs number and suppressive function. Tregs cells express a unique transcription factor, encoded by the X-linked gene *Foxp3*, responsible for maintaining the phenotype of Tregs. Loss of Foxp3 can cause fatal autoimmune diseases, such as IPEX syndrome (Immune dysregulation, Polyendocrinopathy, Enteropathy, X-lined) in humans, which is also responsible for scurfy mice’s lethal phenotype (Fontenot et al., 2003; Shohei Hori, 2003)

The Foxp3 protein comprises multiple domains, including a N-terminal proline-enriched region, a leucine zipper related to dimer formation, a zinc finger, and a C-terminal FKH DNA-binding domain. Dimerization of Foxp3 via the leucine zipper is required for binding to the GTAAACA motif via the C-terminal forkhead (FKH) domain. Studies on IPEX patients have provided many critical functional insights into Foxp3 protein; large numbers of IPEX are missense mutations localized at the FKH domain, and some break Foxp3 dimerization. Thus, Foxp3 must bind the DNA target to fulfill its role undoubtedly. Paradoxically, the full-length of Foxp3 can not bind to DNA unless its N-terminal 181 amino acids ablation. These studies suggested that this N-terminal of Foxp3 could prevent its DNA binding in an autoinhibitory fashion (Koh et al., 2009; Guo et al., 2014). Whether this conformation change of Foxp3 could affect Tregs function is unknown.

Although multiple exons encode the N-terminal of Foxp3, exon 2 has attracted much of our attention. In human but not mouse Tregs, Foxp3 has two main isoforms: full-length Foxp3 and an alternatively spliced form in which exon 2 is deleted (Foxp3Δ2). The exact molecular mechanism of the alternative splicing of Foxp3 pre-mRNA is unknown. However, TCR activation can significantly promote Foxp3Δ2 formation. Although more than 70% of Foxp3 is made up of Foxp3Δ2 in the activated Tregs, the functional role of Foxp3Δ2 in Tregs remains unclear. *In vitro* studies have found that ectopic expression of Foxp3Δ2 in human CD4^+^CD25^+^CD45RA^+^ T cells by retroviral infection can induce a partial Tregs phenotype and confers some immunosuppressive functions (Allan et al., 2005). Expression of both Foxp3 isoforms simultaneously would likely result in a more robust suppressive function.

According to their origin, Tregs are classified into thymus-derived Tregs (tTregs) and peripherally induced Tregs (pTregs). tTregs originate from the thymus, while pTregs originate from peripheral naïve CD4^+^ T cells, especially in the mucosal tissue. tTregs and pTregs have different TCR repertoire and function in distinct immunological contexts. The transcription factor Helios and the surface molecule neuropilin1 are usually used as markers to differentiate tTregs from pTregs (Thornton et al., 2010; Weiss et al., 2012; Yadav et al., 2012). Deleting the conserved nucleotide sequence (CNS1) of Foxp3 results in a defect in pTregs selection, with the loss of Tregs in the intestines and eventual development of colitis (Ye Zheng, 2010; Josefowicz et al., 2012; Schlenner et al., 2012). It is generally believed that the development and differentiation of pTregs are stimulated by low doses of exogenous antigens, such as intestinal microorganisms and food, and that pTregs develop in a microenvironment marked by the presence of TGF-β, RA, IL-2, and other co-stimulators. Microbial products such as short-chain fatty acids facilitate the differentiation of pTregs (Arpaia et al., 2013; Atarashi et al., 2013; Furusawa et al., 2013; Patrick M. Smith, 2013). pTregs recognizing commensal antigens are enriched in the colon (Lathrop et al., 2011; Nutsch et al., 2016) and can further induce the expression of RORγt, which is a master regulator of Th17 cells. RORγt^+^ Tregs constitute the main population of colonic Tregs whose deficiency has been linked to microbial dysbiosis, increased inflammatory Th17 cells, and greater susceptibility to colitis in various mouse models (Ohnmacht et al., 2015; Sefik et al., 2015; Ye et al., 2017; Xu et al., 2018; Al Nabhani et al., 2019; Neumann et al., 2019). Interestingly, RORγt and Foxp3 also directly interact, which is mediated by the LXXLL-containing region of exon 2, and this interaction of RORγt with Foxp3 suppresses RORγt-mediated IL-17A promoter activation (Du et al., 2008; Zhou et al., 2008). The consequence of the interruption of Foxp3-RORγt in pTregs has not been investigated.

The present study was undertaken to determine whether interruption of the autoinhibitory conformation of Foxp3 could affect Tregs function. By using AlphaFold2 structure predictions and *in vitro* experiments, we demonstrate that the N-terminal domain of Foxp3 inhibited its DNA binding by masking the FKH domain. Exon 2 ablation results in a colossal conformation change allowing the FKH domain to bind to its target DNA. Furthermore, we report that sustaining the DNA binding activity of Foxp3 negatively impacts tTregs function by the inhibit expression of BATF and CD25, leading to a decrease in Foxp3Δ2 tTregs activation and function. As a result, this study reveals that the auto-inhibitory loop of Foxp3 is necessary for optimal tTregs activation and function. On the other hand, removing exon 2 has little effect on peripherally induced Tregs and even positively regulated homeostasis of RORγt^+^ pTregs due to the interruption of the interaction of Foxp3-RORγt. Therefore, splicing of exon 2 deferentially regulates tTregs and pTregs homeostasis, which might provide a unique switch for human Tregs in response to various environmental clues.

## RESULTS

### Deletion of exon 2 of Foxp3 results in constitutive DNA binding

Recently, deep-learning methods such as AlphaFold2 (Tunyasuvunakool et al., 2021) and RoseTTAFold (Minkyung Baek, 2021) can extract information from the extensive database of known protein structures in the Protein Data Bank (PDB). Thus, these deep-learning methods outperform more traditional approaches that explicitly model the folding process. Here, we used AlphaFold2 to analyze the structure of Foxp3 and another dominant splicing isoform Foxp3Δ2. The predicted image revealed that two isoforms of Foxp3 had a very different structure. The full-length Foxp3 protein resembles a sandwich based on AlphaFold2, the FKH domain, and a zinc finger-leucine zipper in the middle covered by two pieces of N terminal regions on both sides. The exon 2 region functions as a pin that ties the N-terminal structure with the FKH and zinc finger-leucine zipper. Deletion of exon 2 freed the FKH domain of the leucine zipper and allowed them to be exposed on the outside (Figure 1A, B). The further comparison of the structure of the predicted Foxp3Δ2 protein with the resolved FKH domain structure with DNA (Bandukwala et al., 2011) supports the notion that the structure of full-length Foxp3 restrains DNA binding. However, the deletion of exon 2 causes a substantial conformation change that releases the FKH and leucine zipper domain that enables DNA binding (Figure 1C).

**Figure 1.**
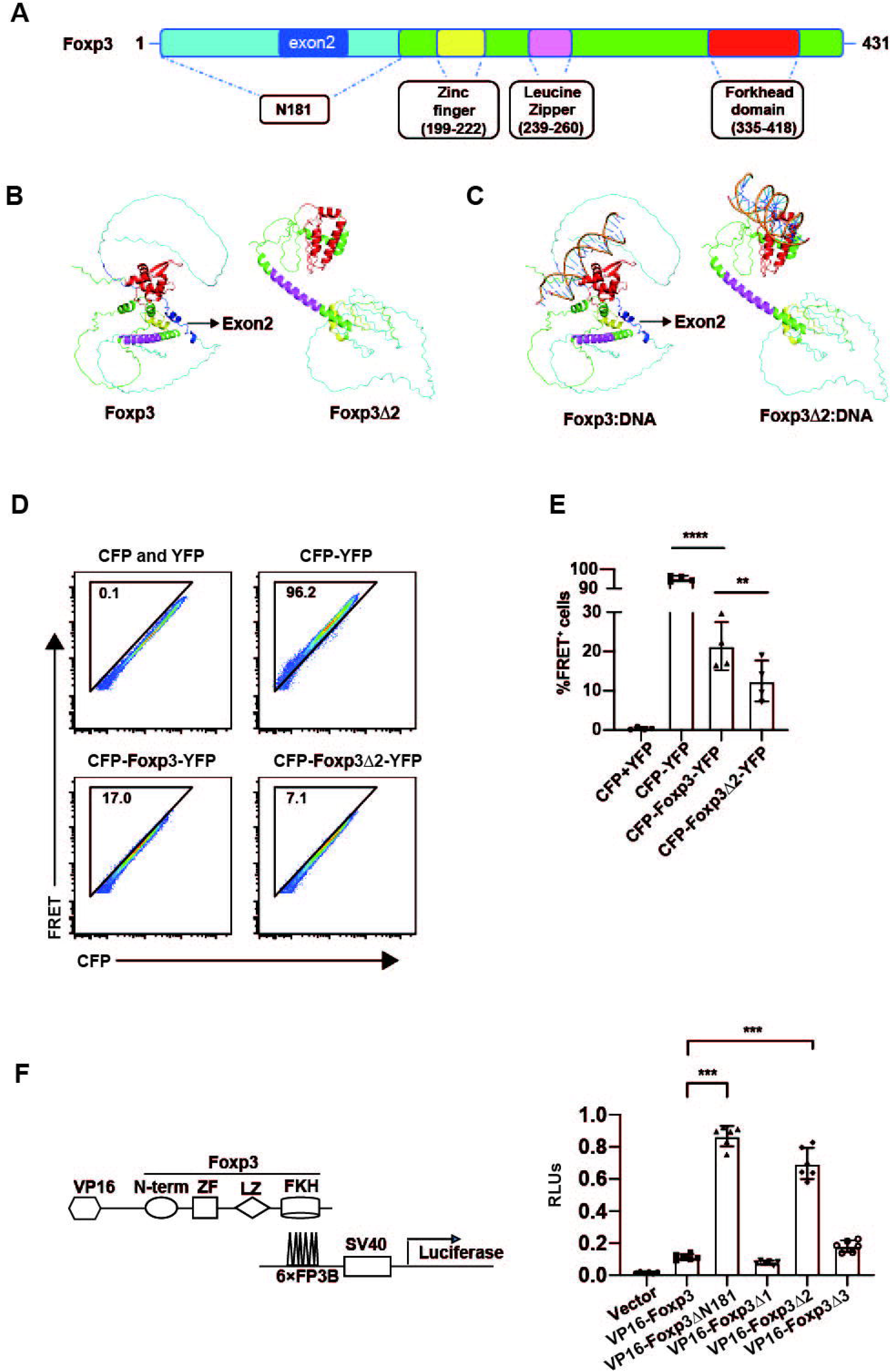
Deletion of exon 2 of Foxp3 results in constitutive DNA binding. (**A**) The functional domains of Foxp3. Domains are represented by different colors: N181 (cyan), exon 2 (blue), zinc finger (yellow), leucine zipper (magenta), FKH (red), others (green). (**B**) Protein structure of Foxp3 and Foxp3Δ2 based on AlphaFold2. The colors of the different domains in the structure correspond to (A). (**C**) Foxp3 and targeted DNA based on AlphaFold2. (**D**) Flow cytometry analysis of the percentage of FRET-positive cells after overexpression of Foxp3 or Foxp3Δ2 fused with CFP at the N-terminus and YFP at the C-terminus. CFP and YFP overexpressed 293T cells were negative control, CFP-YFP overexpressed 293T cells were positive control. (**E**) Quantification of the results in (D). (**F**) Analysis of the DNA-binding capability of Foxp3 or its mutants by comparing the ratio of firefly luciferase to renilla luciferase. The data represent the mean ± SD. The results are representative of at least three independent experiments with n ≥ 3. ns, no significance, *P < 0.05, **P < 0.01, ***P < 0.001, unpaired t test.

To verify this structure model, we performed experiments using a FRET assay, in which the fluorescent proteins CFP and YFP were fused to the N-terminus and C-terminus, respectively, of Foxp3 and Foxp3Δ2. If the N-terminal domain of Foxp3 folds toward the C-terminal FKH domain, the CFP fused to the N-terminal domain will approach the YFP fused to the C-terminal domain. When the CFP is excited by a laser to emit fluorescence, resonance transfer of the fluorescence energy will stimulate the adjacent YFP to emit fluorescence. We detected the FRET signal by expressing the CYP-Foxp3-YFP fusion protein in 293 cells, by contrast, which was significantly decreased by expressing the CFP-Foxp3Δ2-YFP fusion (Figure 1D, E). These results supported the structure model from AlphaFold2 prediction that the N-terminal domain of the full-length Foxp3 protein folds toward the C-terminal FKH domain.

To evaluate the DNA-binding activity of Foxp3 directly, we used a luciferase-based reporter assay (Foxp3Luc), in which a VP16 domain was fused to the N-terminus of different Foxp3 mutants. Upon VP16-Foxp3 binding to the target site (6XFoxp3 binding GTAAACA motif), the VP16 domain can recruit numerous transcriptional activators to start the robust transcription of the mini SV40 promoter-driven luciferase gene (Guo et al., 2014). We first verified that the binding activity of full-length Foxp3 to the target DNA was poor, and deleting the N-terminal 181 amino acids of Foxp3 enhanced its DNA-binding activity remarkably. Consistent with the FRET assay, the DNA-binding activity of Foxp3, indicated by the luciferase activity, was almost wholly retrieved when exon 2 was deleted. In contrast, there was no significant change compared with the entire length of Foxp3 when exon 1 or exon 3 was deleted (Figure 1F). Together, our results demonstrated that alternative splicing of human Foxp3Δ2 broke the inhibitory loop and generated constitutive DNA-binding activity.

### Auto-inhibitory loop of Foxp3 is necessary for optimal Tregs function

To gain functional insight into Foxp3Δ2, we generated exon 2-deficient mice using CRISPR/Cas9-based genome editing (Figure S1A). We speculated a sequential splicing event would allow Foxp3Δ2 protein to express in mouse Tregs once ablation of exon 2. Two mouse strains, *Foxp3^Δ2/Y^* and *Foxp3^mut/Y^* mice with 13 bases missing in exon 2 were obtained after micro-injection. We first verified the successful knockout of exon 2 at both DNA and mRNA levels. PCR from genomic DNA and RT-PCR from RNA demonstrated that exon 2 was removed in the *Foxp3^Δ2/Y^* mice. The sequencing result from RT-PCR products revealed indeed splice exon 1 to exon 3 (Figure S1B, C). High throughput RNA-seq analysis with purified Tregs from the *Foxp3^Δ2/Y^* mice further supported this notion (Figure S1D). Consequently, we checked the expression of Foxp3Δ2 at the protein level by using various antibodies that recognized different epitopes of Foxp3. As expected, we found mAb NRRF-30 targeting the epitope 1-75 and a polyclonal Ab targeting the epitope 250-350 were still reactive with the Foxp3Δ2 either by intracellular staining or western blot, but not FJK-16S, which is a mAb recognizing amino acids 75-125 within exon 2 (Figure S1E, F). These results further validate that exon 1 is linked to exon 3 and that the resulting protein was complete except for the absence of the 35 amino acids encoded by exon 2. Thus, the deletion of exon 2 indeed led to generating a mouse stain that only expresses Foxp3Δ2 isoform in Tregs.

Unlike the *Foxp3^mut/Y^* mice, which have a frameshift mutation in exon 2, and developed spontaneous and fatal scurfy-like autoimmune diseases, *Foxp3^Δ2/Y^* male mice were not abnormal compared with WT male mice in the same litter in terms of appearance and size (Figure S2A-C). Serum auto-antibody detection by ELISA revealed no significant difference in anti-dsDNA levels in *Foxp3^Δ2/Y^* male mice compared with WT male mice in the same litter (Figure S2C). In addition, we examined whether *Foxp3^Δ2/Y^* mice are more resistant to experimental autoimmune encephalomyelitis (EAE), a model for multiple sclerosis (MS). In *Foxp3^WT/Y^* and *Foxp3^Δ2/Y^* mice immunized with MOG, no detectable differences in the time of disease onset and peak score of the disease were observed (Figure S2D). Interestingly, further analysis revealed dysfunction of Tregs in the *Foxp3^Δ2/Y^* male mice. Compared with WT mice, the proportion of CD44^+^CD62L^-^ cells among CD4^+^Foxp3^-^ (Tconv) and CD8^+^ cells was higher in *Foxp3^Δ2/Y^* mice, and levels of the inflammatory cytokines IFN-γ and IL-4 were also increased in the spleen and lymph nodes (Figure 2A-D). We also found that the proportion of Tregs among total CD4^+^ T cells was significantly higher in *Foxp3^Δ2/Y^* male mice than in WT mice (Figure 2E, F), consistent with the inflammatory environment. Moreover, in a mouse metastatic tumor model established by intravenous injection of B16 melanoma cells, the number of lung metastases was significantly reduced in *Foxp3^Δ2/Y^* male mice (Figure 2G). These results suggest activating self-reactivated CD4 and CD8 T cells strengthened their anti-tumor ability. The above results indicate that although breaking the auto-inhibitory loop of Foxp3 enhances the DNA binding activity, sustaining the DNA binding activity of Foxp3 could negatively impact Tregs function.

**Figure 2.**
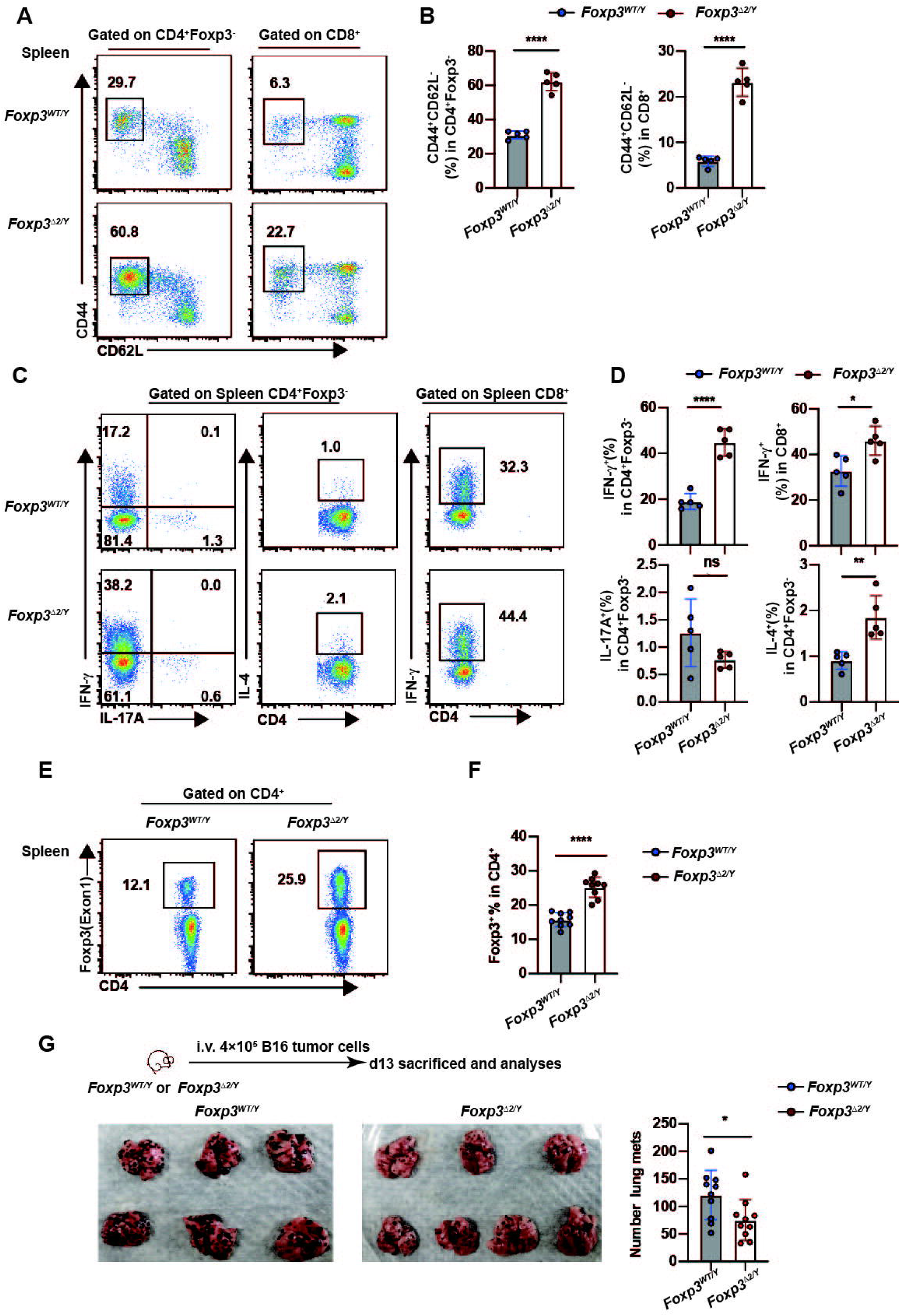
Auto-inhibitory loop of Foxp3 is necessary for optimal Tregs function. (**A-B**) Flow cytometry analysis of CD44 and CD62L expression in CD4^+^Foxp3^-^ and CD8^+^ T cells (A) and frequency of CD44^+^CD62L^-^ cells (B) from the spleen. (**C-D**) Cells were stimulated with PMA and ionomycin for 3 h before intracellular staining. Flow cytometry analysis of IL-17A, IFN-γ and IL-4 expression in CD4^+^Foxp3^-^ T cells and IFN-γ expression in CD8^+^ T cells from the spleen (C) and frequency of IL-17A^+^, IFN-γ^+^ and IL-4^+^ in CD4^+^Foxp3^-^ T cells and frequency of IFN-γ^+^ in CD8^+^ T cells from the spleen (D). (**E-F**) Flow cytometry analysis of the expression of Foxp3 in the spleen of *Foxp3^WT/Y^* and *Foxp3^Δ2/Y^* mice (E). The proportion of Tregs is shown (F). (**G**) 4 × 10^5^ cells B16 cells were i.v. injected into *Foxp3^WT/Y^* and *Foxp3^Δ2/Y^* mice. Melanoma metastases in the lungs were excised (left) and counted after approximately 13 days (right). The data represent the mean ± SD. The results are representative of at least three independent experiments with n ≥ 3. ns, no significance, *P < 0.05, **P < 0.01, ***P < 0.001, unpaired t test.

### Foxp3 isoforms differentially regulate tTregs and pTregs homeostasis

To accurately analyze the phenotype of Foxp3Δ2 Tregs under physiological conditions, we used *Foxp3*^*WT*/Δ*2*^ heterozygous female mice for further investigation. The Foxp3 gene is on the X chromosome, and one allele is randomly inactivated in the female mice. Thus theoretically, both WT and Foxp3Δ2 Tregs would exist in the same *Foxp3*^*WT*/Δ*2*^ heterozygous female mice as a one-to-one ratio. We found that in the thymus, lymph nodes, spleen, liver, and lungs, the proportion of Foxp3Δ2 Tregs among CD4^+^ T cells was much lower than that of WT Tregs (Figure 3 A, B), although the expression level of Foxp3 in Foxp3Δ2 Tregs, indicated as Mean Fluorescent Intensity (MFI), was not decreased (Fig 3C). This result demonstrated that Foxp3Δ2 Tregs had a competitive disadvantage over the full-length Foxp3 Tregs in most peripheral tissue. This deficiency provides a logical explanation for the auto-reactive phenotype found in male *Foxp3^Δ2/Y^* mice.

**Figure 3.**
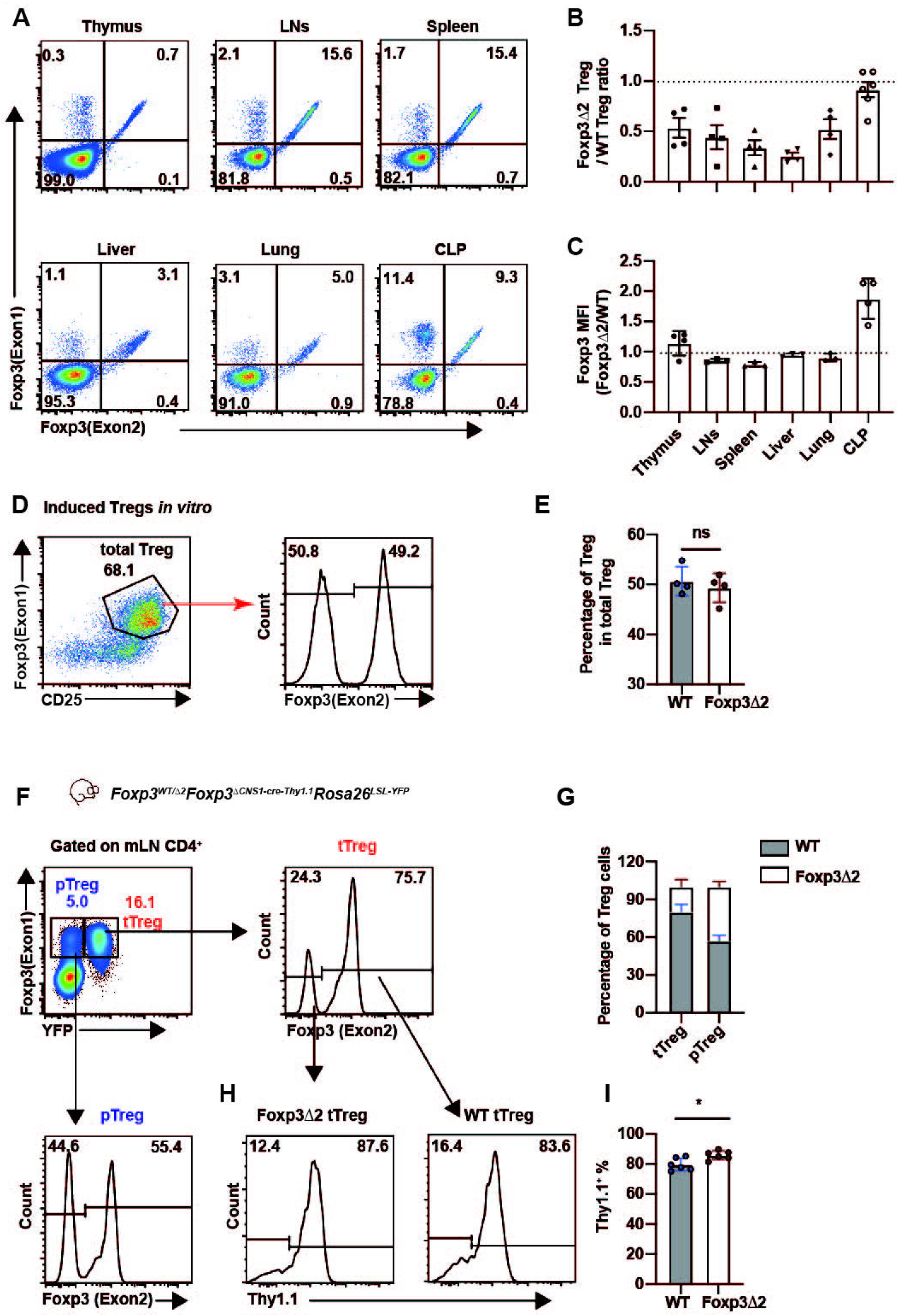
Foxp3 isoforms differentially regulate tTregs and pTregs homeostasis. (**A**) Analysis of the percentage of Foxp3Δ2 Tregs and WT Tregs in CD4^+^ T cells in the thymus, lymph nodes, spleen, liver, lungs and cLP of *Foxp3*^*WT*/Δ*2*^ female mice. (**B**) Quantification of the results in (A). (**C**) The Foxp3 MFI of Foxp3Δ2 Tregs and WT Tregs in CD4^+^ T cells in the thymus, lymph nodes, spleen, liver, lungs and cLP of *Foxp3*^*WT*/Δ*2*^ female mice. (**D**) CD4^+^ cells were enriched from *Foxp3*^*WT*/Δ*2*^ female mice and cultured in plates coated with anti-CD3 (0.5 μg/ml) and anti-CD28 (1 μg/ml) in the presence/absence of TGFβ (2 ng/ml), IL-2 (200 U/ml), and RA (25 nM) for 4 d. Foxp3 and CD25 expression were then analyzed by flow cytometry. (**E**) The percentages of Foxp3Δ2 Tregs and WT Tregs in iTregs. (**F**) Gated on CD4^+^ T cells from mesenteric lymph nodes (mLN) of *Foxp3^RFP/Δ2^Foxp3^ΔCNS1-Thy1.1^Rosa26^LSL-YFP^* female mice. The percentages of Foxp3Δ2 Tregs and WT Tregs in tTregs and pTregs were analyzed. (**G**) Quantification of the data in (F). (**H**) The expression of Thy1.1 in Foxp3Δ2 tTregs and WT tTregs. (I) Quantification of the data in (H). The data represent the mean ± SD. The results are representative of at least three independent experiments with n ≥ 3. ns, no significance, *P < 0.05, **P < 0.01, ***P < 0.001, paired t test.

The only exception to this defect is in the gut; we found no difference in the percentages of these two groups of cells in the colonic lamina propria (cLP). Given that pTregs were highly enriched in colonic lamina propria (cLP), we speculated Foxp3Δ2 might have a distinct function in tTregs and extrathymically generated pTregs. We first tested whether the loss of exon 2 affects the formation of induced Tregs (iTregs) *in vitro*. Naïve CD4^+^CD25^-^ cells from *Foxp3*^*WT*/Δ*2*^ heterozygous female mice were stimulated *in vitro* with antibodies against CD3 and CD28 in the presence of TGF-β, RA, and IL-2 for four days. We found the percentage of Foxp3Δ2^+^ and Foxp3 full-length iTregs were almost identical, suggesting exon 2 of Foxp3 is dispensable for iTregs induction (Figure 3D, E). To further explore this possibility, we crossed the Foxp3Δ2 mice with a tTregs fate-mapping mouse line *Foxp3^ΔCNS1-cre-Thy1.1^Rosa26^LSL-YFP^*, in which only the mature tTregs express the Thy1.1-Cre fusion protein that YFP can label(Zhang et al., 2017). Thy1.1 expression also provides a convenient indicator to evaluate tTregs stability, as termination of Thy1.1 expression precedes the loss of Foxp3. In the *Foxp3*^*WT*/Δ*2*^*Foxp3^ΔCNS1-cre-Thy1.1^Rosa26^LSL-YFP^* heterozygous female mice, YFP^+^ gated bona fide tTregs population possessed less than 25% Foxp3Δ2 Tregs (Figure 3F, G). Moreover, this competitive disadvantage of Foxp3Δ2 Tregs was not due to the instability of Foxp3Δ2, as the expression of Thy1.1 showed that Foxp3Δ2 tTregs were less likely to lose the expression of Thy1.1 than WT Tregs, suggesting that Foxp3Δ2 Tregs were even more stable (Figure 3H, I). In contrast, we found that in Foxp3^+^YFP^-^ pTregs, Foxp3Δ2 Tregs almost have the same ratio as the WT Tregs (Figure 3F, G). Together, these results demonstrated that Foxp3Δ2 selectively regulates the homeostasis of thymus-derived tTregs.

### Foxp3Δ2 Tregs have a defect in activation

To gain detailed mechanistic insight into Foxp3Δ2 tTregs, we performed RNA-seq analysis of Tregs isolated from *Foxp3^RFP/Δ2^Foxp3^ΔCNS1-cre-Thy1.1^Rosa26^LSL-YFP^* female mice. CD4^+^YFP^+^RFP^+^ (WT Tregs) and CD4^+^YFP^+^RFP^-^ (Foxp3Δ2 Tregs) cells were sorted from lymph nodes and spleen to a typical purity of >95%, and RNA-seq and bioinformatics analyses were conducted. In total, we detected 638 differentially expressed genes (DEGs, FDR < 0.05) between Foxp3Δ2 and WT Tregs (Figure 4A). Gene ontology (GO) analysis indicated that the DEGs were most enriched in genes related to T cell activation (Figure 4B). Notably, many genes involved in Tregs activation and differentiation, such as *Il1rl1* (encoding the IL-33 receptor ST2), *Tigit, Icos, Klrg1, Tnfrsf18, Cd44*, and *Ctla4*, were downregulated in Foxp3Δ2 Tregs. In contrast, genes associated with the naive or resting state, including *Ccr7, Sell, Tcf7* (encoding TCF1), *Klf2*, and *Satb1*, were upregulated (Figure S3). The gene set enrichment analysis (GSEA) using a previously published signature of effector Tregs (eTregs) (Dias et al., 2017) further confirmed that genes downregulated during eTregs differentiation were enriched in Foxp3Δ2 Tregs. However, genes upregulated during eTregs differentiation were enriched in WT Tregs (Figure 4C). Moreover, Flow cytometry analysis further verified these differential expressions of various genes (Figure 4D-G). These observations suggested that Foxp3Δ2, breaking the auto-inhibitory loop of Foxp3, acts as a brake to stop tTregs activation and differentiation.

**Figure 4.**
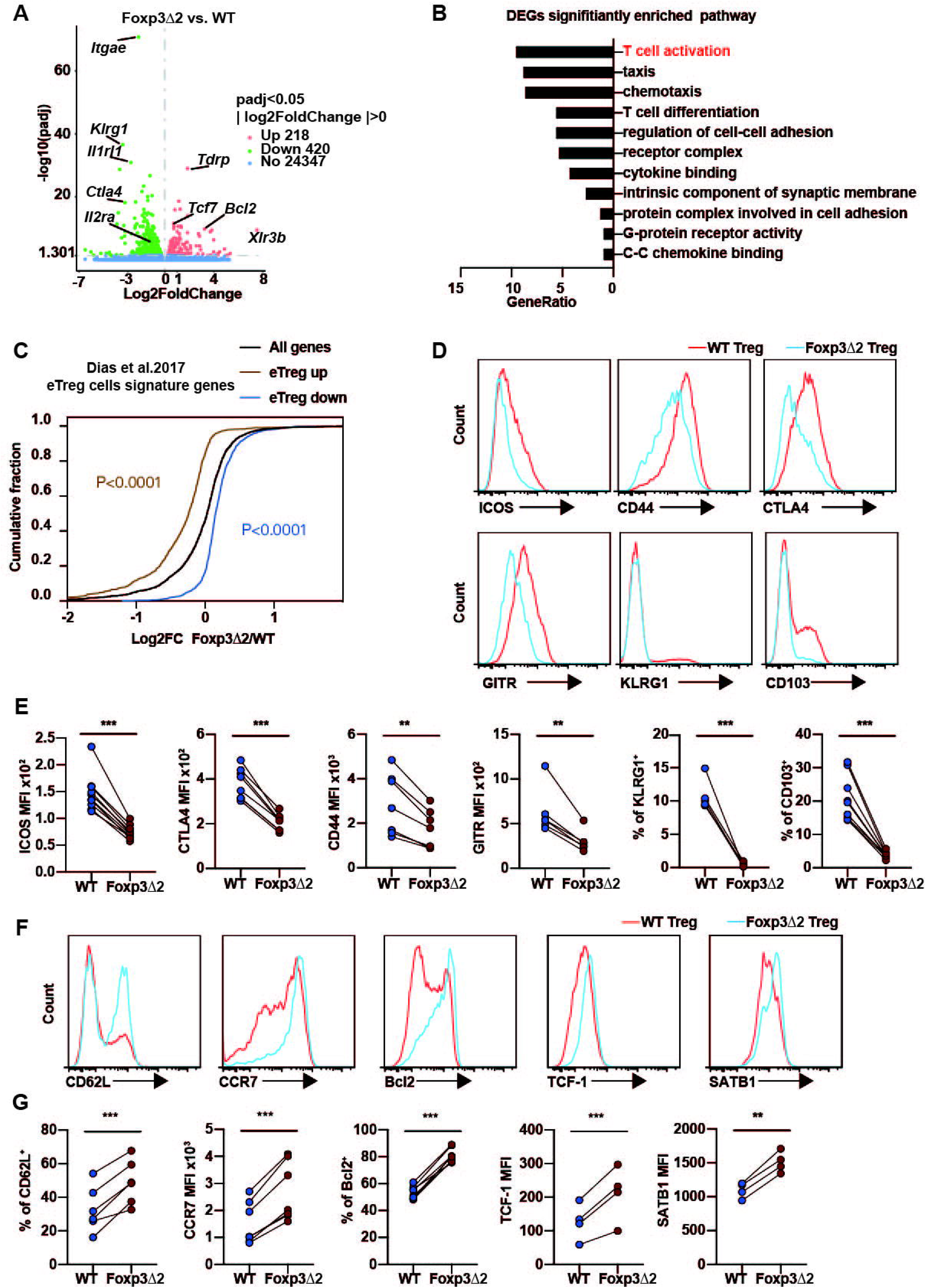
Foxp3Δ2 Tregs lack multiple activation markers and effector molecules. (**A**) FACS-sorted CD4^+^YFP^+^RFP^+^ (WT tTregs) and CD4^+^YFP^+^RFP^-^ (Foxp3Δ2 tTregs) from the spleen and lymph nodes of *Foxp3^RF3/Δ2^Foxp3^ΔCNS1-Thy1.1^ Rosa26^LSL-YFP^* female mice were analyzed by RNA-seq. Genes that were upregulated (218 genes) and downregulated (420 genes) in Foxp3Δ2 Tregs are shown in the volcano map. (**B**) Top 11 significantly enriched pathway of differentially expressed genes (DEGs). (**C**) Empirical cumulative distribution function plots showing the gene signatures of effector Tregs (eTreg) cells. Two-sided Kolmogorov–Smirnov test. (**D**) Representative histograms of ICOS, CD44, CTLA4, GITR, KLRG1 and CD103 in splenic Foxp3Δ2 Tregs and WT Tregs from *Foxp3*^*WT*/Δ*2*^ female mice. (**E**) MFIs and percentages of the indicated genes in each population as (D) shown. (**F**) Representative histograms of CD62L, CCR7, Bcl2, TCF1 and SATB1 in splenic Foxp3Δ2 Tregs and WT Tregs from *Foxp3^*WT*/Δ*2*^* female mice. (**G**) MFIs and percentages of the indicated genes in each population as (F) shown. The data represent the mean ± SD. The results are representative of at least three independent experiments with n ≥ 3. ns, no significance, *P < 0.05, **P < 0.01, ***P < 0.001, paired t test.

### Foxp3Δ2 Tregs are less sensitive to IL-2

Tregs activation and suppression function are controlled by two major signalizing pathways, IL-2 and TCR. Previous RNA-seq analysis also showed that *Il2ra*, a high-affinity receptor for IL-2, was significantly lower in Foxp3Δ2 Tregs than in WT Tregs (Figure S3A). In addition, GSEA revealed that the IL-2 signaling pathway genes were enriched in WT Tregs (Figure 5A). Flow cytometry analysis further showed that the protein expression of CD25 was significantly lower in Foxp3Δ2 Tregs than in WT Tegs in the thymus, spleen, and lymph nodes of *Foxp3*^*WT*/Δ*2*^ female mice (Figure 5B, C). Consistent with these results, we found that the phosphorylation level of STAT5 in an *in vitro* IL-2 stimulation test was significantly lower in Foxp3Δ2 Tregs than in WT Tregs when stimulated with a low concentration of IL-2. This difference in STAT5 phosphorylation can be bypassed by a high concentration of IL-2 (Figure 5D, E). The above results showed that Foxp3Δ2 Tregs are less sensitive than WT Tregs to IL-2 due to their low expression of high-affinity receptors to IL-2, which is likely responsible for the homeostasis defect of Foxp3Δ2 Tregs in the most peripheral tissue.

**Figure 5.**
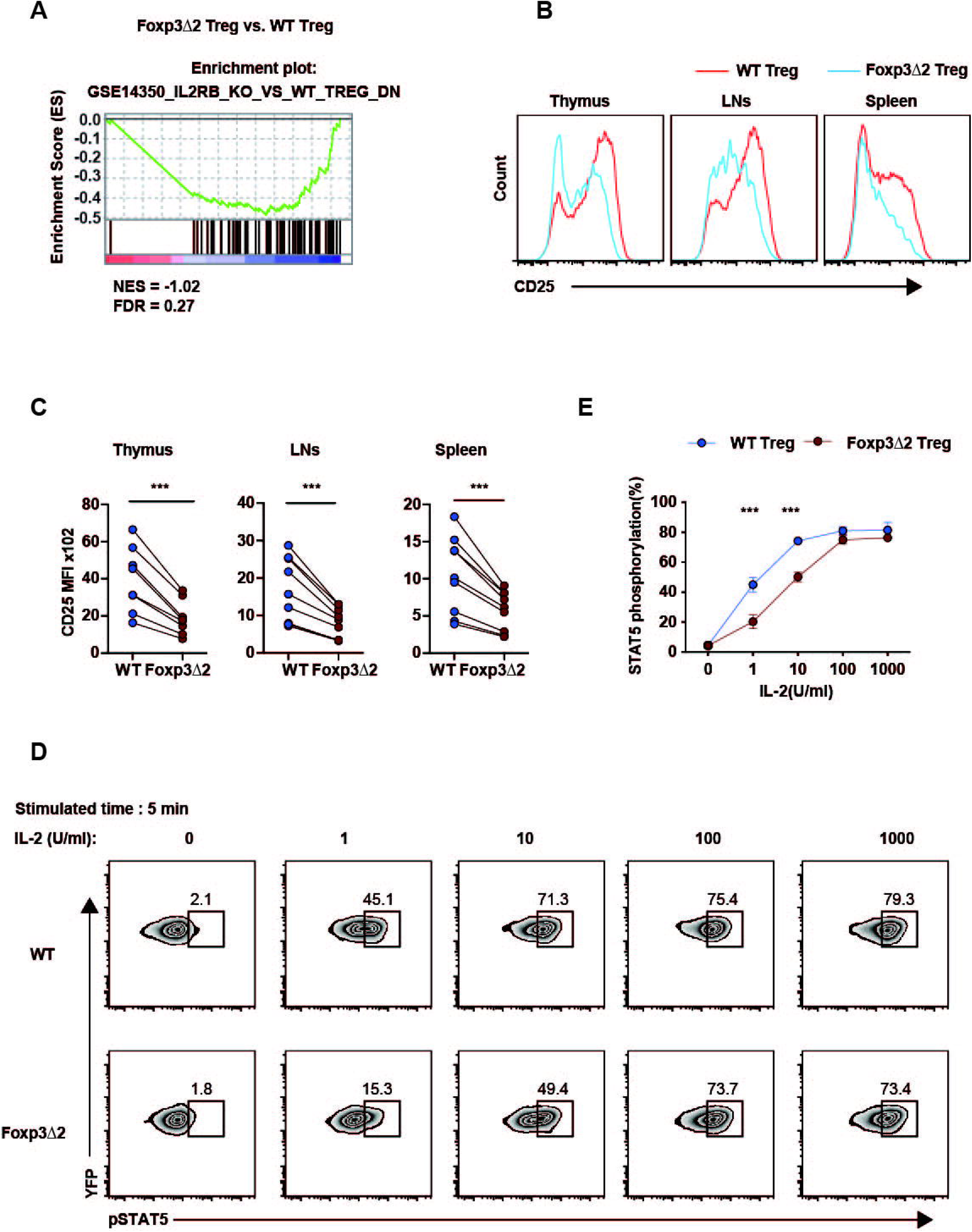
Foxp3Δ2 Tregs are less sensitive to local IL-2. (**A**) GSEA showing the enrichment of IL-2 pathway-related genes in Foxp3Δ2 Tregs. (**B-C**) Flow cytometry analysis of the expression of CD25 in thymus, LN, and spleen of Foxp3Δ2 Tregs and WT Tregs from *Foxp3^WT/Δ2^* female mice. (**D**) Phosphorylation of STAT5 at Try694 (pSTAT5 [Y694]) cells stimulated with 0, 1, 100, 1000 U/mL rhIL-2 for 5 min at 37°C in WT and Foxp3Δ2 Tregs from *Foxp3^WT/Y^Foxp3^GFP-cre^Rosa26^LSL-YFP^* and *Foxp3^Δ2/Y^Foxp3^Cre^Rosa26^LSL-YFP^* male mice, respectively. (**E**) Quantification of STAT5 phosphorylation in Foxp3Δ2 and WT Tregs. The data represent the mean ± SD. The results are representative of at least three independent experiments with n ≥ 3. ns, no significance, *P < 0.05, **P < 0.01, ***P < 0.001, paired t test.

### Enhanced DNA-binding activity contributes to decreased BATF expression in Foxp3Δ2 Tregs

Aside from IL-2 signaling, TCR signaling is another crucial pathway in controlling Tregs activation and function (Levine et al., 2014). We next asked whether the naive phenotype of Foxp3Δ2 Tregs is due to an inability to produce TCR complex and initiate TCR signaling. We found that TCRβ, and ZAP-70 expression in Foxp3Δ2 Tregs was comparable to that in WT Tregs (Figure 6A, B). In addition, the level of CD5, a T cell-marker that is quantitatively correlated with proximal TCR signal intensity, was similarly high in Foxp3Δ2 and WT Tregs, indicating that the deficiency of activation of Foxp3Δ2 Tregs is not due to the defect of proximal TCR signaling (Figure 6A, B). Furthermore, we evaluated the phosphorylation level of S6, a downstream target of mTOR, which is activated by TCR signaling and is essential for the homeostasis and function of Tregs (Zeng et al., 2013). The flow cytometry results showed that the phosphorylation level of S6 did not differ significantly between Foxp3Δ2 and WT Tregs stimulated by the same TCR signal *in vitro* (Figure 6 C, D).

**Figure 6.**
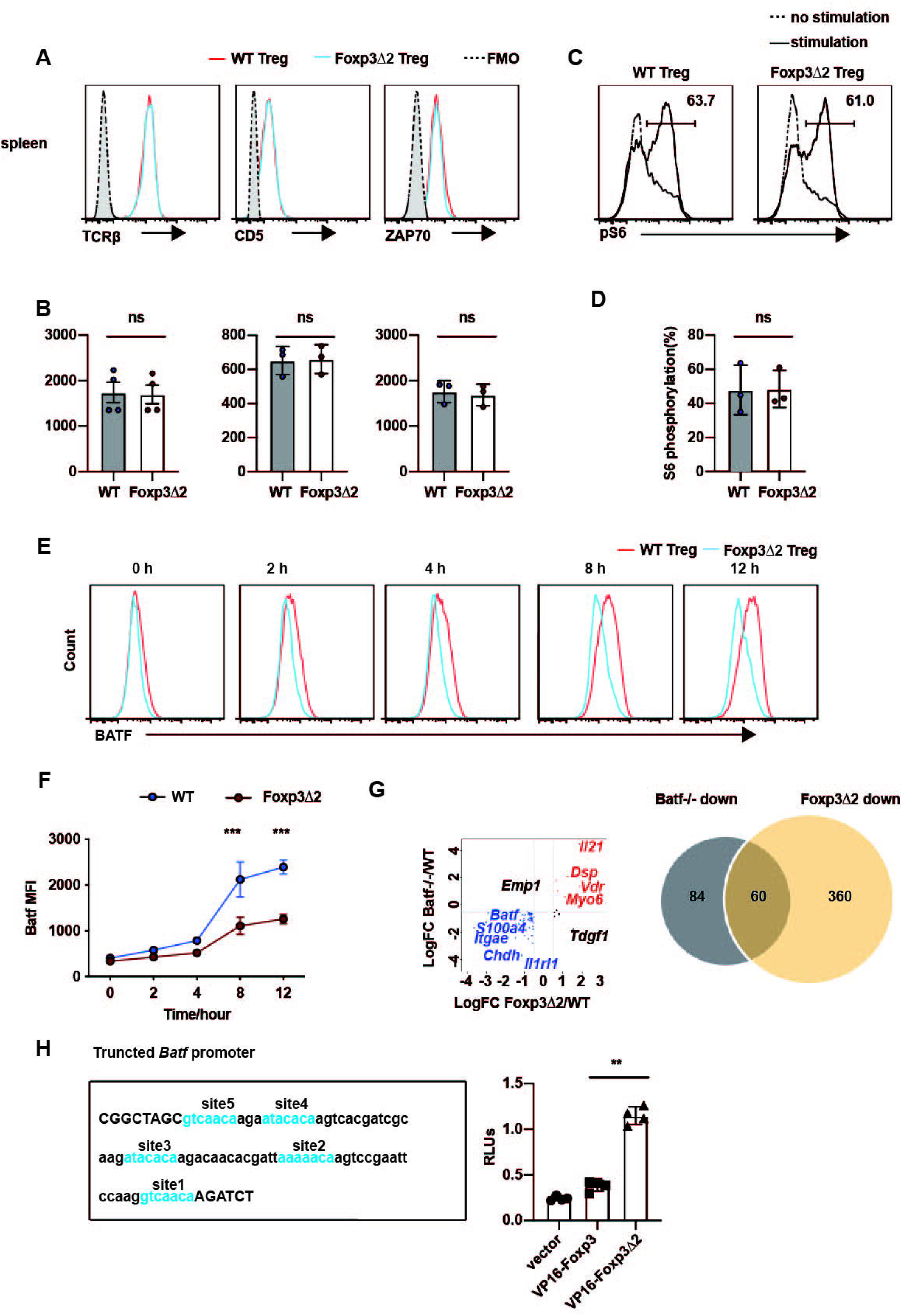
Decreased BATF expression in Foxp3Δ2 Tregs contributes to tTregs homeostasis. (**A**) Representative histograms of TCRβ, CD5 and ZAP70 expression in the spleen and lymph nodes of *Foxp3*^*WT*/Δ*2*^ female mice. (**B**) MFIs of the indicated genes in each population. (**C-D**) Representative histograms of S6 phosphorylation in WT and Foxp3Δ2 Tregs from *Foxp3^WT/Y^Foxp3^GFP-cre^Rosa26^LSL-YFP^* and *Foxp3^Δ2/Y^Foxp3^Cre^Rosa26^LSL-YFP^* male mice, respectively. (**E**) Flow cytometry analysis of the expression of BATF in Tregs from *Foxp3*^*WT*/Δ*2*^ heterozygous female mice after stimulation with anti-CD3 and anti-CD28 antibodies *in vitro* for different time periods. (**F**) Quantification of the results in (E). (**G**) Overlap between genes differentially expressed in the Foxp3Δ2 Tregs vs. WT comparison and those previously identified as *Batf-/-* Tregs DEGs. (**H**) Analysis of the DNA-binding capability of Foxp3 or Foxp3Δ2 to the *Batf* promoter (left) by comparing the ratio of firefly luciferase to renilla luciferase. The data represent the mean ± SD. The results are representative of at least three independent experiments with n ≥ 3. ns, no significance, *P < 0.05, **P < 0.01, ***P < 0.001, paired t test.

AP-1 transcription factors such as IRF4 and BATF, induced by TCR signaling, are critical for Tregs activation and function (Zheng et al., 2009; Hayatsu et al., 2017). Interestingly, the previous study has also shown that an IPEX mutation, Foxp3^A384T^, binds the *Batf* promoter more strongly than WT Foxp3, inhibits the expression of BATF, and eventually reduces the activation level of Tregs (Hayatsu et al., 2017). Although the expression of BATF was slightly downregulated in Foxp3Δ2 Tregs compared with WT Tregs in the spleen and lymph nodes, Foxp3Δ2 Tregs was indeed poorly responsive to TCR when the two groups of Tregs received the same strength of TCR stimulation *in vitro*. The difference in BATF expression between Foxp3Δ2 and WT Tregs increased significantly in a time-dependent manner (Figure 6E, F). Moreover, RNA-seq analysis revealed that the genes with lower expression levels in Foxp3Δ2 Tregs overlapped significantly with those with lower expression levels in Batf^-/-^ Tregs. (Figure 6G). The *Batf* promoter contains five FKH motif-like sequences, and previous data suggest WT Foxp3 was merely bound to two GTCAACA and two ATACACA in the EMSA assay. We cloned these five FKH motif-like sequences into the pGL3 vector and co-transfected them with VP16-Foxp3 or VP16-Foxp3Δ2 in HEK293T cells. We observed that *Batf* promoter binding by VP16-Foxp3Δ2 was approximately 3-fold higher than that by VP16-Foxp3 (Figure 6H). Therefore, we concluded that the open conformation of Foxp3Δ2 facilitates its binding to *Batf* promoter and decreases BATF expression, down-regulated their sensitivity to TCR signal.

### The expression of RORγt-related genes is increased in Foxp3Δ2 Tregs in the large intestine

Given the fact that exon 2 deletion can also interrupt Foxp3-RORγt protein-protein interaction, we next focused on the RORγt^+^ Tregs, which constitute the about 60% colonic pTregs cells in B6 mice that differentiate locally in response to bacterial antigens from 15–20 days of age onward (Ohnmacht et al., 2015; Sefik et al., 2015). Although our initial experiment found that the proportions of Foxp3Δ2 Tregs and WT Tregs were not significantly different in the cLP in *Foxp3*^*WT*/Δ*2*^ heterozygous female mice, further RNA-seq analysis did reveal some unique features of Foxp3Δ2 Tregs in the gut. In total, we detected 1029 DEGs (FDR < 0.05) between Foxp3Δ2 and WT Tregs (Figure S4A). GSEA revealed that the IL-17 signaling pathway was dramatically enriched in Foxp3Δ2 Tregs (Figure S4B), increasing *IL-17, Il23r, Ccl20, Il1r1*, *Vdr, Nr1d1*, and *F2rl2*, but downregulation of *Il1rl1, Klrg1, Ikzf2, Gata3* and *Lrrc32* in Foxp3Δ2 Tregs (Figure 7A). We also analyzed intracellular flow cytometry to validate these results at a single cell level. More strikingly, we found that Foxp3Δ2 Tregs were overwhelming RORγt^+^ and uniformly expressed a high level of c-Maf, an upstream transcriptional factor that plays a substantial role in the differentiation and function of RORγt^+^ pTregs (Figure 7B, C). In addition, Foxp3Δ2 Tregs had low Helios, ST2, and Gata3, which are highly expressed in colonic tTregs (Figure 7B, C). In addition, we found that Foxp3Δ2 Tregs in the colonic lamina propria had high expression of IL-17A, the target gene downstream of RORγt, and low expression of GPR15, the gene suppressed by RORγt, compared with WT Tregs (Fig7 B, C).

**Figure 7:**
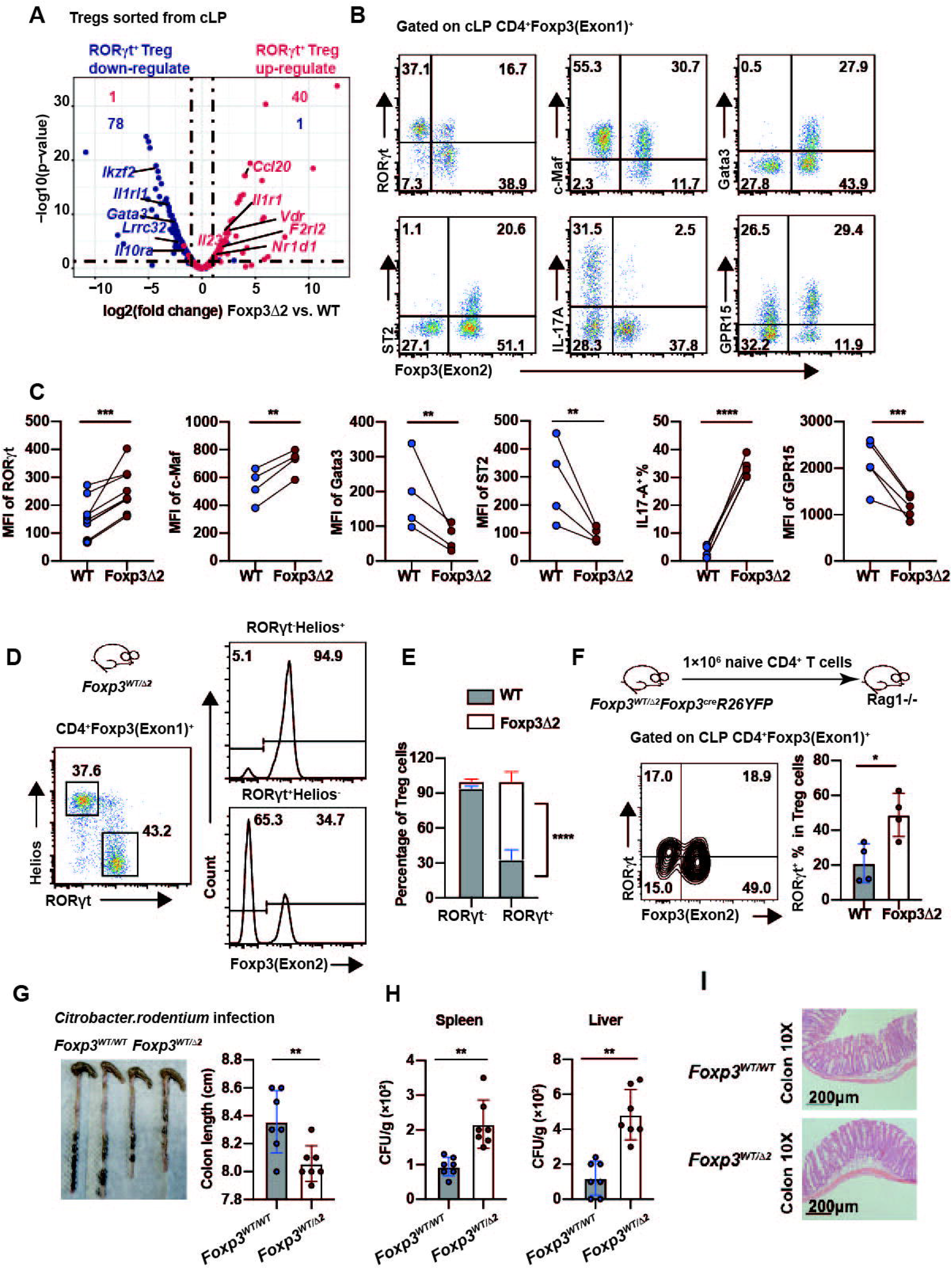
Enhanced expression of RORγt in Foxp3Δ2 Tregs increases the susceptibility of *Foxp3*^*WT*/Δ*2*^ mice to *C. rodentium* infection. (**A**) Distribution of upregulated and downregulated RORγt^+^ Tregs signature genes among DEGs. (**B-C**) The expression of RORγt, c-Maf, ST2 and Gata3 in WT and Foxp3Δ2 Tregs in the colonic lamina propria (cLP) in *Foxp3*^*WT*/Δ*2*^ female mice was analyzed by flow cytometry. The MFI of each marker in the indicated population is shown (C). cLP lymphocytes were stimulated with PMA, ionomycin, and monensin for 3–4 h and examined for Foxp3 and IL-17A expression. The expression of GPR15 was also analyzed in the absence of stimulation. The frequency of IL-17A^+^ and MFI of GPR15 in the indicated population are shown (C). (**D-E**) The expression of RORγt in WT and Foxp3Δ2 Tregs from the cLP of *Foxp3*^*WT*/Δ*2*^ female mice fed by C57BL/6 or Balb/c mothers was analyzed by flow cytometry. The frequency of RORγt^+^ in the indicated population is shown (E). (**F-G**) *Foxp3*^*WT*/Δ*2*^ and *Foxp3^WT/WT^* female mice were infected with 10^9^ CFU of *C. rodentium* at 6-8 weeks of age. (F) Intestine length after approximately 2 weeks; (G) *C. rodentium* counts in the spleen and liver on day 13. The data represent the mean ± SD. The results are representative of at least three independent experiments with n ≥ 3. ns, no significance, *P < 0.05, **P < 0.01, ***P < 0.001, paired t test.

Helios expressing tTregs and RORγ^+^ expressing pTregs, are two phenotypically and functionally different subpopulations in the colon. Therefore, We re-analyzed flow data by first gating on RORγ^+^ and Helios^+^ respectively. Consistent with the early finding in the second lymphoid organ, Foxp3Δ2 tTregs poorly competed with WT tTregs, and this defect is more severe in the cLP; in fact, over 95% Helios^+^ population were WT Tregs. In sharp contrast, among colonic RORγ^+^ pTregs, Foxp3Δ2 Tregs clearly have a competitive advantage over WT RORγ^+^ pTregs (Figure 7D, E). Thus, expression of the multiple transcriptional signatures of RORγ^+^ Tregs in the colon Foxp3Δ2 tTregs due to combined effects of loss of Helios^+^ tTregs and accumulation of RORγ^+^ pTregs. Given the fact of tTregs can regulate RORγt under certain circumstances, to rule out this possibility, we directly transferred naive CD4^+^Foxp3^-^ Tconv cells from *Foxp3^GFP-cre^Foxp3*^*WT*/Δ*2*^ mice female to the RAG KO mice for four weeks, and bona fide pTregs were checked. Indeed, we detected that Foxp3Δ2 Tregs have a much more significant percentage of RORγt^+^ cells than the WT control (Figure 7F). RORγt^+^ induction in Foxp3Δ2 Tregs similarly relied on bacterial antigens, which can be limited by maternal IgA, as RORγt^+^ Foxp3Δ2 cells from *Foxp3*^*WT*/Δ*2*^ mice were significantly decreased by either treating with antibiotics or fostering with Balb/c mother at birth (Figure S5 A-E).

Finally, to study the specific functions of Foxp3Δ2 RORγt^+^ pTregs, we used a model of intestinal infection in which the mouse pathogen *Citrobacter rodentium* was used to infect adult *Foxp3*^*WT*/Δ*2*^ or *Foxp3^WT/WT^* female mice. We speculated that WT tTregs in the mosaic *Foxp3*^*WT*/Δ*2*^ female mice would compensate for the defect of Foxp3Δ2 tTregs, while Foxp3Δ2 RORγ^+^ pTregs would provide superior immune suppression effects in the colon. Indeed, compared with *Foxp3^WT/WT^* mice, *Foxp3*^*WT*/Δ*2*^ mice had more severe colitis by both the appearance of the gut and H&E staining, correlated with increased bacterial translocation to extraintestinal sites, including the spleen and liver, and decreased Th17 response in the cLP of *Foxp3*^*WT*/Δ*2*^ mice (Figure 7G-I, Figure S6). Thus, interruption of the Foxp3-RORγt interaction in the Foxp3Δ2 RORγt^+^ pTregs benefits them for better adapting to the gut environmental conditions, which was correlated with a dampened antibacterial inflammatory response.

## DISCUSSION

The present study documented that the auto-inhibitory loop of Foxp3 is necessary for optimal tTregs homeostasis, activation, and function. By releasing the inhibition of TCR-induced BATF induction, and expressing a high level of CD25, Foxp3 WT tTregs are more sensitive to extra-cellar antigens and IL-2 stimulation, thus having a competitive advantage over the Foxp3Δ2 tTregs. We also documented that exon2 deletion has little impact on iTregs and *in vivo* peripheral induced pTregs, and even confers a competitive advantage over the WT once expressed RORγt in pTregs. Given that alternative splicing is a highly dynamic event in response to various environment clues, we speculated that alternative splicing could provide a unique switch for human Tregs regulation.

The DNA-binding ability of full-length Foxp3 *in vitro* is fragile and is influenced by the N-terminal region of the protein, given the fact that N-terminal proline-enriched region is intrinsically disordered and has resisted structural determination. We modeled the Foxp3 structure by using AlphaFold2, which can extract information from the extensive database of known protein structures. We found that the N-terminal region of Foxp3 impedes the binding of the FKH domain of Foxp3 to DNA. Exon 2 is a central component of the auto-inhibitory loop, whose deletion results in a colossal conformation change allowing the FKH domain to bind to its target DNA. How full-length Foxp3 binds to target genes is still under determination. Foxp3 was modulated by various post-translational modifications, including phosphorylation, acetylation, ubiquitylation, and methylation. It also interacts with a large number of transcription factors and forms multiprotein complexes of 400–800 kDa or larger. We speculated that post-translational modifications and/or protein and protein interaction might provide the chance to open the auto-inhibitory structure.

Surprisingly, sustaining the DNA binding activity of Foxp3 negatively impacts tTregs function by decreasing Foxp3Δ2 tTregs activation and function. We observed that Foxp3Δ2 tTregs had low expression levels of many activations- and effector-related proteins, such as ICOS, KLRG, CTLA4, and CD44, and high expression levels of naïve Treg-related proteins, such as TCF1, SATB1, and CD62L. In addition, Foxp3Δ2 tTregs are less sensitive to extra-cellar antigens and IL-2 stimulation. The mouse phenotype of Foxp3Δ2 Tregs shares similarity with a human IPEX patient who had a deletion of base 305 in the exon2 of Foxp3, which caused frameshift mutations of full-length of Foxp3 but had no effect on Foxp3Δ2 formation. In this patient, although Foxp3Δ2 protein was intact and the proportion of Tregs was even higher than that found in healthy people, the IPEX patient had mild autoimmunity, and the effector T cells in were in a more active state. Moreover, an identical competitive disadvantage of Foxp3Δ2 tTregs was an event in the healthy carrier mother. Therefore, we concluded that in both mouse and human Tregs, the auto-inhibitory loop of Foxp3 is necessary for optimal tTregs homeostasis, activation, and function. Mechanistically, we revealed that the defect of Foxp3Δ2 tTregs is primarily due to sustained foxp3 binding to *Batf*, which is a direct target gene of Foxp3 and plays an essential role in Tregs activation and function. We found that the expression level of BATF was significantly lower in Foxp3Δ2 Tregs than in Foxp3 Tregs when the Tregs were stimulated with the same concentration of CD3/CD28 antibody. Moreover, genes with lower expression levels in Foxp3Δ2 Tregs overlapped significantly with those with lower expression levels in Batf-/- Tregs. Interestingly, another IPEX-associated mutation Foxp3 A384T, can also specifically target BATF; whether A384T mutation would affect the conformation of Foxp3 need further investigation.

We also documented that exon 2 deletion can regulate tTregs and pTregs distinctly. In the colon form mosaic *Foxp3*^*WT*/Δ*2*^ female mice, we found that over 95% tTregs were WT Tregs, but in sharp contrast, among colonic RORγ^+^ pTregs, a significant bias toward Foxp3Δ2 Tregs. Foxp3 forms large complexes with other transcription factors or modifying enzymes to regulate gene expression. The LXXLL-containing region of exon 2 has been reported to bind the AF2 regions of RORγt and suppresses RORγt-mediated IL-17A activation. On the other hand, mice with RORγt deficiency in Foxp3-expressing Tregs develop severe oxazolone-induced colitis and trinitrobenzene-sulfonic acid-induced colitis (Ohnmacht et al., 2015; Sefik et al., 2015). Our result demonstrated that interruption of the Foxp3-RORγt interaction will release the inhibition effect of Foxp3, which benefits Foxp3Δ2 RORγt^+^ pTregs for better adapting to the gut environmental conditions.

In conclusion, the present study provides a new perspective on a dominant human isoform, Foxp3Δ2. Our results against the notion that Foxp3Δ2 is a useless product of alternative splicing, we propose Foxp3Δ2 is a unique switch that balances human tTregs and pTregs.

## Supporting information

immunity_v1

## Acknowledgments

We are particularly grateful to the Pathogenic Microbiology and Immunology Public Technology Service Center for its support. We gratefully acknowledge Dr. Mingzhao Zhu and Dr. Zhaolin Hua for suggestions and all members of our laboratory for discussions.

## Author Contributions

Q. G., X. Z., J. G., W. Z., W. X., J. Z. performed experiments. X. Zhou conceived and supervised the project. F.Z. and B.H. helped with some experiments. X. Z. and Q. G. wrote the manuscript.

## STAR ⍰ Methods

### Key resources table

| REAGENT or RESOURCE | SOURCE | IDENTIFIER |
| --- | --- | --- |
| CD4-PerCP-Cy5.5 (RM4-5) | eBioscience | Cat#45-0042-82 |
| CD8α-AF700 (53-6.7) | BioLegend | Cat#100730 |
| CD45R/B220-PE (RA3-6B2) | BioLegend | Cat#103208 |
| CD44-APC (IM7) | eBioscience | Cat#17-0441-83 |
| CD62L-PE (MEL-14) | BioLegend | Cat#104408 |
| CD25-APC (PC61.5) | eBioscience | Cat#17-0251-82 |
| KLRG1-APC (2F1) | eBioscience | Cat#17-5893-82 |
| ICOS-APC (7E.17G9) | eBioscience | Cat#17-9942-82 |
| CD69-APC (H1.2F3) | eBioscience | Cat#17-0691-82 |
| CD103-APC (2E7) | eBioscience | Cat#17-1031-82 |
| CTLA4-PE (UC10-4B9) | eBioscience | Cat#14-1522-82 |
| Ki67-eF660 (SolA15) | eBioscience | Cat#50-5698-82 |
| CCR7-PE (4B12) | eBioscience | Cat#12-1971-82 |
| GPR15-PE | BioLegend | Cat#373003 |
| Neuropilin-1 (BAF566) | R&D Systems | Cat#BAF566 |
| ST2-Biotin (RMST2-33) | eBioscience | Cat#13-9333-82 |
| TCRβ-APC (H57-597) | eBioscience | Cat#17-5961-81 |
| CD5-APC (53-7.3) | BD Pharmingen | Cat# 550035 |
| Fixable Viability Dye- eFluor™<br>506 | eBioscience | Cat#65-0866-18 |
| CXCR3-APC (CXCR3-173) | eBioscience | Cat#17-1831-80 |
| Brilliant Violet 605™<br>Streptavidin | BioLegend | Cat#405229 |
| Foxp3-BV421 (FJK-16 s) | eBioscience | Cat#48-5773-82 |
| Foxp3-PE (NRRF-30) | eBioscience | Cat#12-4771-82 |
| Foxp3-AF488 (MF-23) | BD Pharmingen | Cat#560403 |
| IFN-γ-PE (XMG1.2) | eBioscience | Cat#12-7311-82 |
| IL-17A-APC (eBio17B7) | eBioscience | Cat#17-7177-81 |
| IL-4-PE (11B11) | eBioscience | Cat#12-7041-82 |
| IL-10-APC (JES5-16E3) | eBioscience | Cat#17-7101-82 |
| pS6RP-S235/236-AF647 (D57.2.2E) | Cell Signaling Technology | Cat# 4851S |
| pSTAT5-PE(47/Stat5(pY694)) | BD Pharmingen | Cat#562077 |
| BATF-PE(S39-1060) | BD Pharmingen | Cat#564503 |
| Helios-PE (22F6) | BioLegend | Cat#137206 |
| TCF1-AF647 (C63D9) | Cell Signaling Technology | Cat#6709S |
| c-Maf-PE (sym0F1) | eBioscience | Cat#12-9855-42 |
| Gata3-PE (TWAJ) | eBioscience | Cat#12-9966-42 |
| Bcl2-AF647 (BCL/10C4) | BioLegend | Cat#633510 |
| RORγt-APC (B2D) | eBioscience | Cat#17-6981-82 |
| RORγt-Bv421 (Q31-378) | BD Pharmingen | Cat#562894 |
| anti-mouse CD3 (145-2C11) | BioXcell | Cat#BE0001-1 |
| anti-mouse CD28 (PV-1) | BioXcell | Cat#BE0015-5 |

### Chemicals, Peptides, and Recombinant Proteins

|  |  |  |
| --- | --- | --- |
| Recombinant human IL-2 | PeptoTech | AF-200-02 |
| Dynabeads Mouse T-Activator CD3/CD28 for T Cell Expansion and Activation | Life Technology | 11456D |
| BioMag goat anti-Mouse IgG | QIAGEN | 310007 |
| BioMag goat anti-Rat IgG | QIAGEN | 310107 |
| BD Phosflow Lyse/Fix buffer (5 ×) | BD Bioscience | 558049 |
| BD Phosflow Perm Buffer III | BD Bioscience | 558050 |
| Superscript III Reverse Transcriptase | Invitrogen | 18080044 |
| UltraSYBG Mixture | CWBIO | 04707494001 |
| RiboLock RNase Inhibitor | Thermo Fisher | CW0957L |
| Protease inhibitor | Roche | 05892791001 |
| Phorbol 12-myristate 13-acetate (PMA) | Sigma | P8139 |
| Ionomycin | Sigma | I0634 |
| Monensin | BioLegend | 420701 |
| RNeasy<sup>®</sup> MicroRNA Kit | QIAGEN | 74004 |
| Foxp3 Staining Buffer Set | eBioscience | 00-5523-00 |
| CellTrace Violet Proliferation Kit | Invitrogen | C34557 |

### Contact for Reagent and Resource Sharing

Further information and requests for resources and reagents should be directed to and will be fulfilled by the Lead Contact, Xuyu Zhou.

## MATERIALS AND METHODS

### AlphaFold2 prediction

Structure predictions of the Foxp3 and Foxp3Δ2 were performed with the AlphaFold2 software following the instructions at the website https://github.com/sokrypton/ColabFold (accessed on 21 November 2021). The final predicted structures were analyzed by PyMOL software.

### Mice

To generate *Foxp3Δ2* mice, the corresponding sgRNA1 and sgRNA2 sequences were designed based on the intron sequences before and after exon 2 of the mouse *Foxp3* gene. The two sgRNAs were cloned into the expression vector px330 carrying the *cas9* gene (Addgene 42230), and then the two px330 plasmids with different sgRNAs were mixed equally at a final concentration of 2.5 ng/μL and injected into fertilized eggs of B6D2F1 × B6 by the microinjection technique. The surviving fertilized eggs were transplanted into the uterus in pseudopregnant female mice, which produced the first generation of *Foxp3Δ2* mice. *Foxp3^GFP-Cre^* (Zhou et al., 2009), *Foxp3^ΔCNS1-cre-Thy1.1^* (Zhang et al., 2017), *Rosa26^LSL-YFP^* (Shankar Srinivas, 2001), and *Foxp3-RFP* knockin mice (Wan and Flavell, 2005) have been described previously. C57BL/6J mice were purchased from HuaFuKang (Beijing). All mice were housed at the specific pathogen-free animal facility at the Institute of Microbiology, Chinese Academy of Sciences. All experiments were conducted according to the guidelines of the Institute of Microbiology, Chinese Academy of Sciences Institutional Animal Care and Use Committee (permit no. 1116050300119).

### Luciferase assay

To determine which exon of the N-terminus affects the DNA-binding ability of Foxp3, we constructed expression plasmids in which exon 1, exon 2 or exon 3 was deleted. Then, HEK293T cells were co-transfected with each of the Foxp3 plasmids and the target plasmid pGL3-NB143, which encodes the tandem Foxp3 binding sequence for use with the dual fluorescence reporting system established in our laboratory (Guo et al., 2014). This system uses Renilla luciferase as an internal standard. Dual luciferase activity was measured using the Dual-Luciferase^®^ reporter assay system (Promega, catalog no. E1960) and a GloMax^®^20/20 luminometer according to the manufacturer’s specifications. Experiments were conducted in triplicate. Data were normalized to control groups. Differences were analyzed for statistical significance using the unpaired Student’s *t* test.

### Lymphocyte isolation

Spleens and lymph nodes were collected from the various mouse lines. Splenocytes were treated with 1 × ACK to lyse red blood cells, and single-cell suspensions were subsequently prepared by mechanical disruption in DMEM supplemented with 2% (v/v) fetal calf serum (FCS). To isolate lymphocytes from the large intestine, the organ was defatted and opened longitudinally, and the luminal contents were removed by vigorous manual shaking of the tissue in PBS. The tissues were then cut into 1–2 cm pieces and incubated in 20 mL of Pre-digest solution [1 × HBSS, plus 1 mM EDTA (Sigma, E4884) and 1 mM DTT (Sigma, D9779) added immediately before use] for 20 min at 37°C with vigorous shaking (250 rpm). Epithelial and immune cells from the epithelial layer were collected by pouring the suspension through a 100-μm strainer (Falcon). The remaining tissue was then incubated in 5 mL of digest solution [200 U/mL Collagenase IV (Worthington, LS004189) and 400 U/mL DNase I (Worthington, LS002139)] for 20 min at 37°C with vigorous shaking (250 rpm). After centrifugation (300 rpm) for 4 min, the supernatant was collected and placed on ice. The undigested tissue was digested for an additional 20 min, passed through a 100 μm strainer, and centrifuged to remove collagenase solution. Lamina propria samples were washed by centrifugation in a 40%:80% Percoll™ gradient (GE, 17-0891-09) to remove debris and enrich for leukocytes, and the lymphocytes in the interphase region were collected.

### Flow cytometry

Antibody staining was performed in ice-cold buffer (RPMI with 2% FCS) for 30 min with dilutions of antibodies. To analyze transcription factors (TFs), cells were fixed, permeabilized, and intracellularly stained for Foxp3, Gata3, RORγt, c-Maf, and Helios according to the manufacturer’s instructions (eBioscience). To analyze the cytokines IL-17A, IFN-γ and IL-4, cells were incubated for 3 to 4 h at 37°C and 5% CO_2_ with 0.5 mM ionomycin, 10 ng/mL PMA, and 3 mM monensin before antibody staining. The assay to detect phosphorylated STAT5 was conducted as previously described (Zhu et al., 2019). Cells were acquired with an LSRII flow cytometer (BD Biosciences), and data were analyzed using FlowJo software.

### Tregs induction *in vitro*

Highly pure naïve CD62L^hi^CD4^+^GFP^-^ T cells from 6- to 8-week-old *Foxp3^GFP-cre^Foxp3*^*WT*/Δ*2*^ mice were sorted by FACS and cultured in plates coated with anti-CD3 (0.5 μg/ml) and anti-CD28 (1 μg/ml) in the presence of TGF-β (2 ng/ml), IL-2 (200 U/ml), and retinoic acid (25 nM) for 4 d. The cells were then analyzed by flow cytometry.

### Western blot

Western blotting was performed to confirm the expression of the Foxp3 protein with deletion of exon 2. CD4^+^ CD25^+^ cells were sorted from *Foxp3^WT/Y^* and *Foxp3^Δ2/Y^* mice, and nuclear proteins were extracted for use in western blotting with an antibody targeting amino acids 250-350 at the C-terminus (Abcam, ab150743) of the Foxp3 protein as previously described. As a positive control, HEK293T cells were co-transfected with plasmids expressing Foxp3 and Foxp3Δ2, and two protein bands of different sizes were differentiated.

### B16 lung metastasis model

B16 melanoma cells were cultured for 3-4 generations and digested with trypsin. The cells were washed once with 1 × PBS and then resuspended in 1 × PBS. Then, 4 × 10^5^ cells in 200 μL were intravenously (i.v.) injected into each *Foxp3^WT/Y^* or *Foxp3^Δ2/Y^* mouse. Thirteen days after cell injection, the mice were sacrificed, the lungs were excised, and melanoma metastases were counted.

### RNA-sequencing

Lymphocytes were isolated from the spleen and lymph nodes of 6- to 8-week-old *Foxp3^RFP/Δ2^Foxp3^ΔCNS1-Thy1.1^R26YFP* female mice and enriched for CD4^+^ T cells using magnetic beads. Then, CD4^+^YFP^+^RFP^+^ (WT Tregs) and CD4^+^YFP^+^RFP^-^ (Foxp3Δ2 tTregs) cells were sorted to a typical purity of >95%. To analyze colon Tregs, Tregs were sorted from the colonic lamina propria of 6- to 8-week-old *Foxp3^GFP-cre^Foxp3^RFP/Δ2^* female mice. CD4^+^GFP^+^RFP^+^ (WT Tregs) and CD4^+^GFP^+^RFP^-^ (Foxp3Δ2 Tregs) cells were sorted to a typical purity of >95%. RNA was extracted using the RNeasy^®^ MicroRNA Kit (QIAGEN 74004). RNA sequencing (RNA-seq) and bioinformatics analysis were conducted by Novogene as described previously (Zhang et al., 2017). Differential expression analysis of the two subsets was performed using the DEGSeq R package (1.20.0). The p values were adjusted using the Benjamini-Hochberg method. A corrected q value of 0.05 and log2 (fold change) of 1 were set as the thresholds for significantly differential expression. The one-tailed Kolmogorov-Smirnov test was used to determine the significance of differences in the distribution of signature genes and all expressed genes.

### Statistical analysis

Comparisons between two experimental groups were performed by Student’s *t* test with a two-tailed distribution. To compare multiple groups, one-way ANOVA was first used to determine whether any of the differences among the means was statistically significant; then, the unpaired Student’s *t* test was used to determine the statistical significance of differences between two specific groups using GraphPad Prism 8.0. Data are presented as means ± SD. P values ≤ 0.05 were considered statistically significant, and asterisks indicate the level of significance as follows: *, P < 0.05; **, P < 0.01; ***, P < 0.001. P values > 0.05 were considered not statistically significant (unmarked or specified as ns).

## Data Availability

Raw RNA-seq data reported in this paper were submitted to the Gene Expression Omnibus database available at https://www.ncbi.nlm.nih.gov/geo/ under the accession code GSE208094 and GSE208370.

## Supplemental information

**Figure S1. Generation and identification of *Foxp3Δ2* mice by CRISPR/Cas9**

(**A**) Schematic diagram: sgRNAs were designed at both ends of exon 2 of the mouse *Foxp3* gene. The underlined sequence is exon 2, and the sequence in red corresponds to the sgRNA. (**B**) Representative genotyping PCR for *Foxp3^WT/Y^*, *Foxp3^Δ2/Y^* and *Foxp3*^*WT*/Δ*2*^ mice. The lower sequence is the Sanger sequencing result, and the deleted sequence includes the complete exon 2 and a partial intron sequence. (**C**) CD4^+^ CD25^+^ cells of *Foxp3^WT/Y^* and *Foxp3^Δ2/Y^* mice were sorted. The RNA was extracted and reverse transcribed into cDNA for use in PCR. The sequencing results below show that exons 1 and 3 are directly linked in the mRNA of *Foxp3^Δ2/Y^* mice. (**D**) RNA-seq analysis of alternative splicing of Foxp3 mRNA from *Foxp3^WT/Y^* and *Foxp3^Δ2/Y^* Tregs. (**E**) Mouse splenocytes were stained with two kinds of Foxp3 antibodies (NRRF-30, which binds exon 1, and FJK-16s, which binds exon 2). (**F**) Nuclear protein was extracted from CD4^+^ CD25^+^ cells sorted from *Foxp3^WT/Y^* and *Foxp3^Δ2/Y^* Tregs and used in western blotting with an antibody targeting the C-terminal 250-350 aa of the Foxp3 protein.

**Figure S2. The phenotype of *Foxp3^Δ2/Y^* mice**

(**A**) Appearance of 8-week-old *Foxp3^WT/Y^* and *Foxp3^Δ2/Y^* mice and 14-day-old *Foxp3^mut/Y^* male mice. (**B**) Spleen and peripheral lymph nodes from the mice shown in (A). (**C**) Body weight and serum anti-dsDNA antibody levels of 5-month-old *Foxp3^WT/Y^* and *Foxp3^Δ2/Y^* male mice. (**D**) The clinical score of *Foxp3^WT/Y^* and *Foxp3^Δ2/Y^* mice after EAE induction. EAE disease progression was monitored daily, and the disease score was assigned on a scale as follows: 0, no disease; 1, loss of tail tonicity; 2, wobbly gait; 3, hindlimb paralysis; 4, hindlimb and forelimb paralysis; 5, moribund.

Each point represents an individual mouse, and the data represent > 3 independent experiments. ns, no significance, *P < 0.05, **P < 0.01, ***P < 0.001.

**Figure S3. RNA-seq analysis of WT and Foxp3Δ2 tTregs**

(**A-B**) The heatmap shows differences in T cell activation-related genes, down-regulated genes (A) and up-regulated genes (B) in Foxp3Δ2 Tregs compared with WT Tregs from RNA-seq results.

**Figure S4. RNA-seq analysis of WT and Foxp3Δ2 Tregs in the cLP**

(**A**) RNA-seq analysis of FACS-sorted CD4^+^GFP^+^RFP^+^ (WT Tregs) and CD4^+^GFP^+^RFP^-^ (Foxp3Δ2 Tregs) from the colonic lamina propria (cLP) of *Foxp3^RFP/Δ2^Foxp3^GFP-cre^* female mice. Genes that were upregulated (532 genes) and downregulated (497 genes) in Foxp3Δ2 Tregs are shown in the volcano map. (**B**) GSEA of the IL-17 signaling pathway in Foxp3Δ2 Tregs.

**Figure S5. Up-regulated RORγt in Foxp3Δ2 Tregs cells depends on microbiota**

(**A**) Technical route of ABX antibiotic treatment of mice: Foxp3WT/Δ2 female mice were fed with sterile water containing 1 mg/ml metronidazole, 0.5 mg/ml vancomycin, 1 mg/ml ampicillin and 1 mg/ml neomycin or sterile water without antibiotics as the control for 4 weeks. (**B**) Flow cytometry analysis showed that the expression of RORγt in Tregs cells of colonic lamina propria in control group and ABX treatment group. (**C**) Statistical analysis the proportion of RORγt^+^ Tregs cells in WT Tregs cells and Foxp3Δ2 Tregs cells between control group and ABX treatment group. (**D**) The genotype of the female offspring was *Foxp3*^*WT*/Δ*2*^ and fed by C57BL/6J or Balb/c mother. The expression of RORγt in Tregs cells of colonic lamina propria of offspring mice fed by C57BL/6J strain and Balb/c strain were analyzed by flow cytometry. (**E**) Proportion of RORγt^+^ Tregs cells in WT Tregs cells and Foxp3Δ2 Tregs cells of the two group.

The data represent the mean ± SD. The results are representative of at least three independent experiments with n ≥ 3. ns, no significance, *P < 0.05, **P < 0.01, ***P < 0.001, paired t test.

**Figure S6. Decreased Th17 response in cLP of *Foxp3*^*WT*/Δ*2*^ mice**

(**A**) After 14 days of *Citrobacter rodentium* infection, the expression of IL-17A in the cLP was detected by flow cytometry. (**B**) The proportion of Th17 cells in CD4^+^ T cells.

The data represent the mean ± SD. The results are representative of at least three independent experiments with n ≥ 3. ns, no significance, *P < 0.05, **P < 0.01, ***P < 0.001, unpaired t test.

