## Supplementary material for "The splicing isoform Foxp3Δ2 releases the autoinhibitory conformation and differentially regulates tTregs and pTregs homeostasis": immunity_v1: Supplemental Figures.pdf

Fig. S1

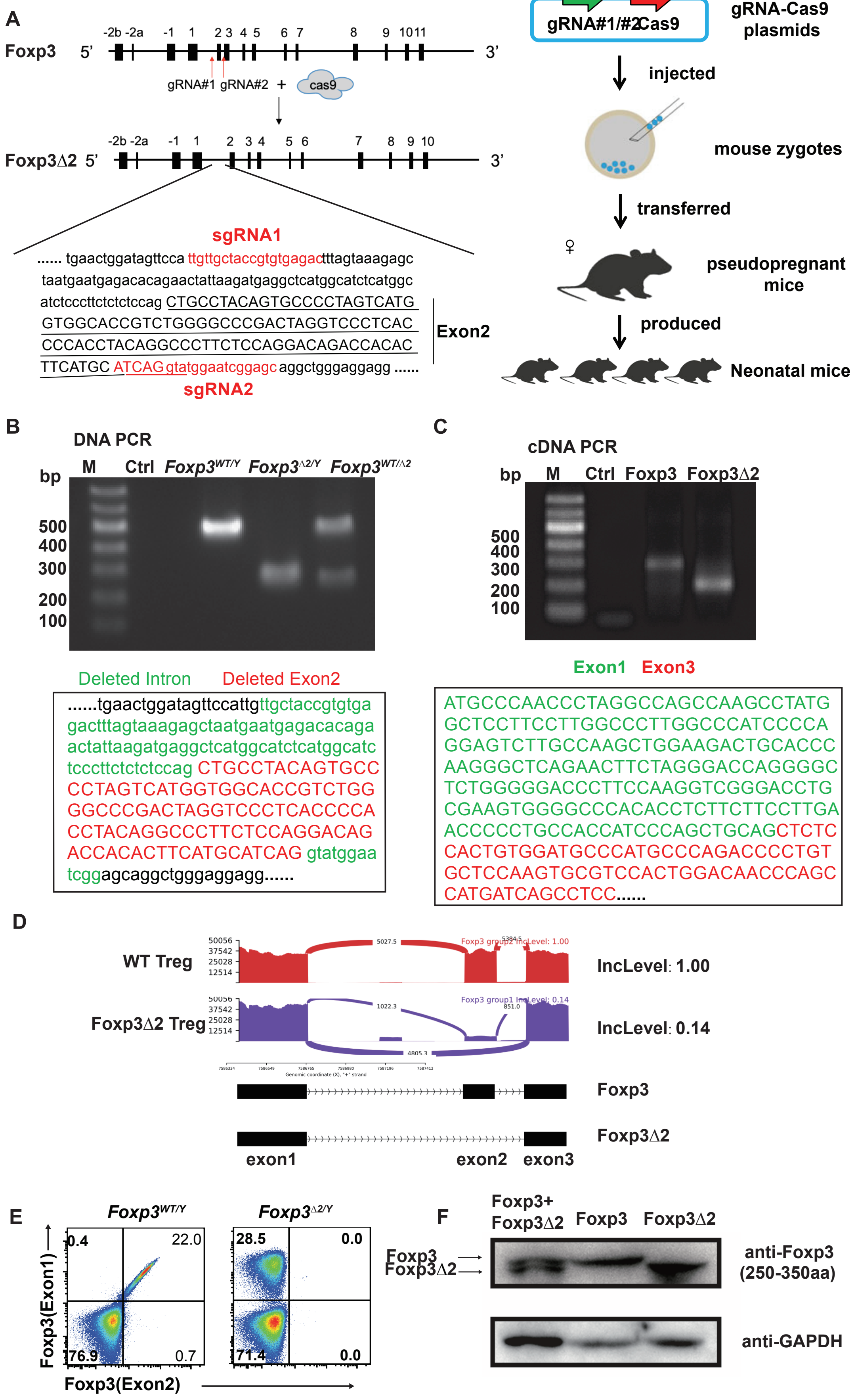
A
Foxp3 5'
-2b -2a -1 1 2 3 4 5 6 7 8 9 10 11
3'
gRNA#1 gRNA#2 + cas9
Foxp3Δ2 5'
-2b -2a -1 1 2 3 4 5 6 7 8 9 10
3'
sgRNA1
...... tgaactggatagttcca ttgttgctaccgtgtgagactttagtaaagagc
taatgaatgagacacagaactattaagatgaggctcatggcatctcatggc
atctcccttctctctccag CTGCCTACAGTGCCCCTAGTCATG
GTGGCACCGTCTGGGGCCCGACTAGGTCCCTCAC
CCCACCTACAGGCCCTTCTCCAGGACAGACCACAC
TTCATGC ATCAG gtatggaatcggagc aggctgggaggagg ......
Exon2
sgRNA2
gRNA#1/#2 Cas9
gRNA-Cas9 plasmids
injected
mouse zygotes
transferred
♀
pseudopregnant mice
produced
Neonatal mice
B
DNA PCR
bp M Ctrl Foxp3WT/Y Foxp3Δ2/Y Foxp3WT/Δ2
500
400
300
200
100
Deleted Intron Deleted Exon2
......tgaactggatagttccattgttgctaccgtgtga
gactttagtaaagagctaatgaatgagacacaga
actattaagatgaggctcatggcatctcatggcatc
tcccttctctctccag CTGCCTACAGTGCC
CCTAGTCATGGTGGCACCGTCTGG
GGCCCGACTAGGTCCCTCACCCCA
CCTACAGGCCCTTCTCCAGGACAG
ACCACACTTCATGCATCAG gtatggaa
tcggagcaggctgggaggagg......
C
cDNA PCR
bp M Ctrl Foxp3 Foxp3Δ2
500
400
300
200
100
Exon1 Exon3
ATGCCCAACCCTAGGCCAGCCAAGCCTATG
GCTCCTTCCTTGGCCCTTGGCCCATCCCCA
GGAGTCTTGCCAAGCTGGAAGACTGCACCC
AAGGGCTCAGAACTTCTAGGGACCAGGGGC
TCTGGGGGACCCTTCCAAGGTCGGGACCTG
CGAAGTGGGGCCCACACCTCTTCTTCCTTGA
ACCCCCTGCCACCATCCCAGCTGCAGCTCTC
CACTGTGGATGCCCATGCCCAGACCCCTGT
GCTCCAAGTGCGTCCACTGGACAACCCAGC
CATGATCAGCCTCC......
D
WT Treg
IncLevel: 1.00
Foxp3Δ2 Treg
IncLevel: 0.14
Foxp3
Foxp3Δ2
exon1 exon2 exon3
E
Foxp3WT/Y
Foxp3Δ2/Y
0.4 22.0 76.9 0.7
28.5 0.0 71.4 0.0
Foxp3(Exon1)
Foxp3(Exon2)
F
Foxp3+ Foxp3Δ2 Foxp3 Foxp3Δ2
Foxp3
Foxp3Δ2
anti-Foxp3 (250-350aa)
anti-GAPDH

**Figure S1. Generation and identification of *Foxp3Δ2* mice by CRISPR/Cas9**

(**A**) Schematic diagram: sgRNAs were designed at both ends of exon 2 of the mouse *Foxp3* gene. The underlined sequence is exon 2, and the sequence in red corresponds to the sgRNA. (**B**) Representative genotyping PCR for $Foxp3^{WT/Y}$, $Foxp3^{\Delta 2/Y}$ and $Foxp3^{WT/\Delta 2}$ mice. The lower sequence is the Sanger sequencing result, and the deleted sequence includes the complete exon 2 and a partial intron sequence. (**C**) $CD4^{+}$ $CD25^{+}$ cells of $Foxp3^{WT/Y}$ and $Foxp3^{\Delta 2/Y}$ mice were sorted. The RNA was extracted and reverse transcribed into cDNA for use in PCR. The sequencing results below show that exons 1 and 3 are directly linked in the mRNA of $Foxp3^{\Delta 2/Y}$ mice. (**D**) RNA-seq analysis of alternative splicing of Foxp3 mRNA from $Foxp3^{WT/Y}$ and $Foxp3^{\Delta 2/Y}$ Tregs. (**E**) Mouse splenocytes were stained with two kinds of Foxp3 antibodies (NRRF-30, which binds exon 1, and FJK-16s, which binds exon 2). (**F**) Nuclear protein was extracted from $CD4^{+}$ $CD25^{+}$ cells sorted from $Foxp3^{WT/Y}$ and $Foxp3^{\Delta 2/Y}$ Tregs and used in western blotting with an antibody targeting the C-terminal 250-350 aa of the Foxp3 protein.

Fig. S2

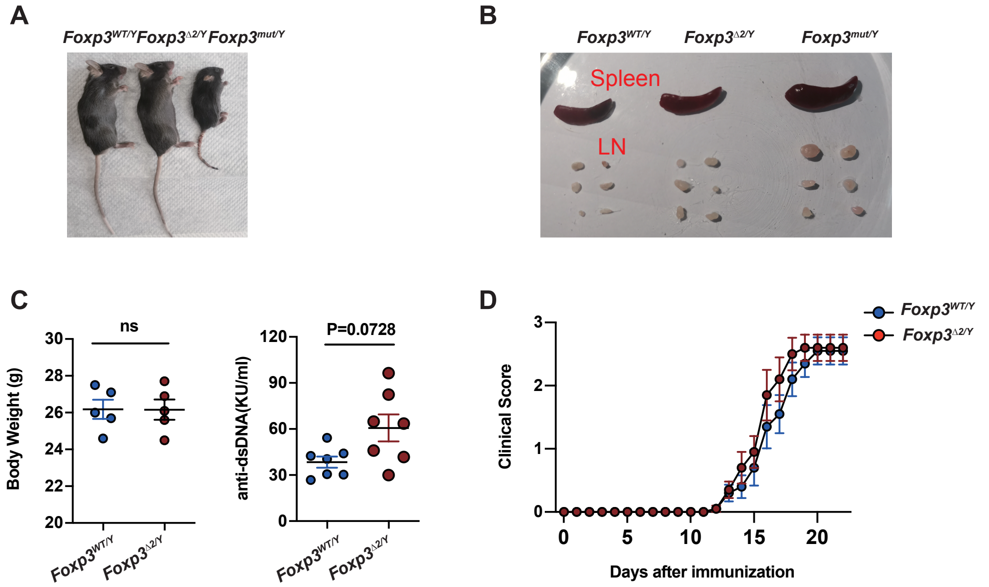
A
Foxp3WT/Y Foxp3Δ2/Y Foxp3mut/Y
B
Foxp3WT/Y
Foxp3Δ2/Y
Foxp3mut/Y
Spleen
LN
C
ns
Body Weight (g)
P=0.0728
anti-dsDNA(KU/ml)
Foxp3WT/Y
Foxp3Δ2/Y
D
Clinical Score
Days after immunization
Foxp3WT/Y
Foxp3Δ2/Y

**Figure S2. The phenotype of Foxp3Δ2/Y mice**

(**A**) Appearance of 8-week-old *Foxp3*$^{WT/Y}$ and *Foxp3*$^{\Delta 2/Y}$ mice and 14-day-old *Foxp3*$^{mut/Y}$ male mice. (**B**) Spleen and peripheral lymph nodes from the mice shown in (A). (**C**) Body weight and serum anti-dsDNA antibody levels of 5-month-old *Foxp3*$^{WT/Y}$ and *Foxp3*$^{\Delta 2/Y}$ male mice. (**D**) The clinical score of *Foxp3*$^{WT/Y}$ and *Foxp3*$^{\Delta 2/Y}$ mice after EAE induction. EAE disease progression was monitored daily, and the disease score was assigned on a scale as follows: 0, no disease; 1, loss of tail tonicity; 2, wobbly gait; 3, hindlimb paralysis; 4, hindlimb and forelimb paralysis; 5, moribund.

Each point represents an individual mouse, and the data represent > 3 independent experiments. ns, no significance, *$P < 0.05$, **$P < 0.01$, ***$P < 0.001$.

Fig. S3

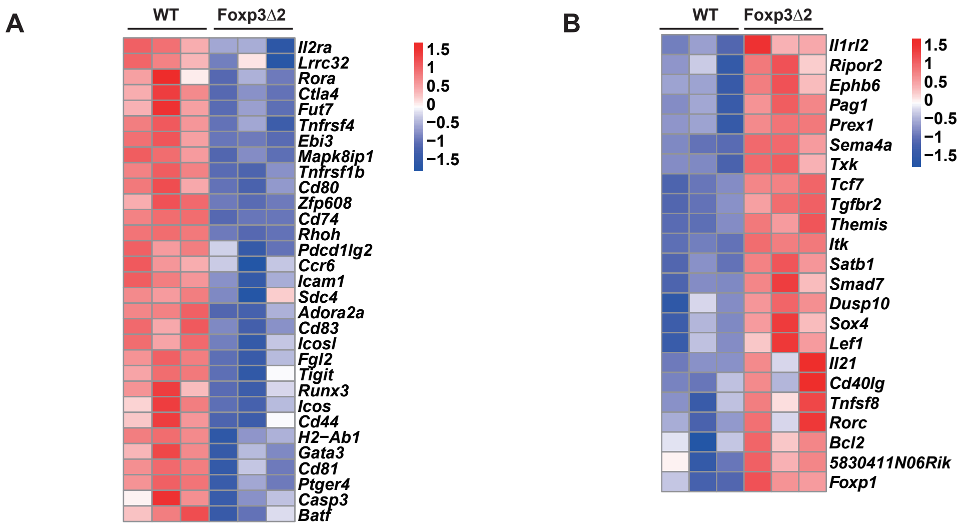
A
WT
Foxp3Δ2
Il2ra
Lrrc32
Rora
Ctla4
Fut7
Tnfrsf4
Ebi3
Mapk8ip1
Tnfrsf1b
Cd80
Zfp608
Cd74
Rhoh
Pdcd1lg2
Ccr6
Icam1
Sdc4
Adora2a
Cd83
Icosl
Fgl2
Tigit
Runx3
Icos
Cd44
H2−Ab1
Gata3
Cd81
Ptger4
Casp3
Batf
1.5
1
0.5
0
−0.5
−1
−1.5
B
WT
Foxp3Δ2
Il1rl2
Ripor2
Ephb6
Pag1
Prex1
Sema4a
Txk
Tcf7
Tgfbr2
Themis
Itk
Satb1
Smad7
Dusp10
Sox4
Lef1
Il21
Cd40lg
Tnfsf8
Rorc
Bcl2
5830411N06Rik
Foxp1
1.5
1
0.5
0
−0.5
−1
−1.5

**Figure S3. RNA-seq analysis of WT and Foxp3Δ2 tTregs**

(**A-B**) The heatmap shows differences in T cell activation-related genes, down-regulated genes (A) and up-regulated genes (B) in Foxp3Δ2 Tregs compared with WT Tregs from RNA-seq results.

### Fig. S4

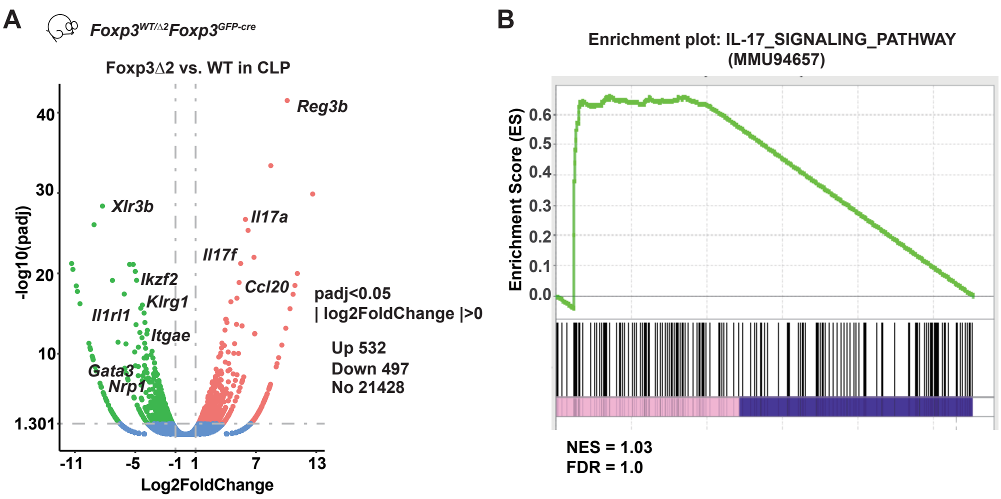

A
Foxp3WT/Δ2Foxp3GFP-cre
Foxp3Δ2 vs. WT in CLP
-log10(padj)
Log2FoldChange
40
30
20
10
1.301
-11
-5
-1
1
7
13
Reg3b
Xlr3b
Il17a
Il17f
Ikzf2
Ccl20
Klrg1
Il1rl1
Itgae
Gata3
Nrp1
padj<0.05
| log2FoldChange |>0
Up 532
Down 497
No 21428
B
Enrichment plot: IL-17_SIGNALING_PATHWAY
(MMU94657)
Enrichment Score (ES)
0.6
0.5
0.4
0.3
0.2
0.1
0.0
NES = 1.03
FDR = 1.0

**Figure S4. RNA-seq analysis of WT and Foxp3Δ2 Tregs in the cLP**

(**A**) RNA-seq analysis of FACS-sorted $CD4^+GFP^+RFP^+$ (WT Tregs) and $CD4^+GFP^+RFP^-$ (Foxp3Δ2 Tregs) from the colonic lamina propria (cLP) of *Foxp3*$^{RFP/\Delta 2}$*Foxp3*$^{GFP\text{-}cre}$ female mice. Genes that were upregulated (532 genes) and downregulated (497 genes) in Foxp3Δ2 Tregs are shown in the volcano map. (**B**) GSEA of the IL-17 signaling pathway in Foxp3Δ2 Tregs.

### Fig. S5

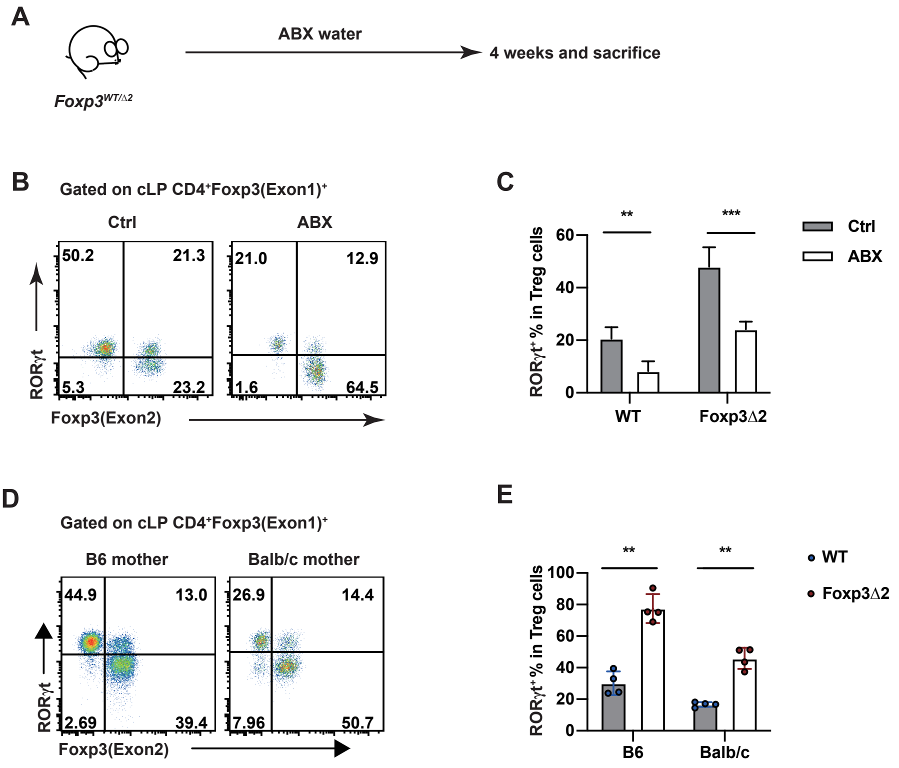
A
ABX water
4 weeks and sacrifice
Foxp3WT/Δ2
B
Gated on cLP CD4+Foxp3(Exon1)+
Ctrl
ABX
50.2
21.3
5.3
23.2
21.0
12.9
1.6
64.5
RORγt
Foxp3(Exon2)
C
RORγt+ % in Treg cells
**
***
Ctrl
ABX
WT
Foxp3Δ2
D
Gated on cLP CD4+Foxp3(Exon1)+
B6 mother
Balb/c mother
44.9
13.0
2.69
39.4
26.9
14.4
7.96
50.7
RORγt
Foxp3(Exon2)
E
RORγt+ % in Treg cells
**
**
WT
Foxp3Δ2
B6
Balb/c

**Figure S5. Up-regulated RORγt in Foxp3Δ2 Tregs cells depends on microbiota**

(**A**) Technical route of ABX antibiotic treatment of mice: Foxp3WT/Δ2 female mice were fed with sterile water containing 1 mg/ml metronidazole, 0.5 mg/ml vancomycin, 1 mg/ml ampicillin and 1 mg/ml neomycin or sterile water without antibiotics as the control for 4 weeks. (**B**) Flow cytometry analysis showed that the expression of RORγt in Tregs cells of colonic lamina propria in control group and ABX treatment group. (**C**) Statistical analysis the proportion of RORγt$^{+}$ Tregs cells in WT Tregs cells and Foxp3Δ2 Tregs cells between control group and ABX treatment group. (**D**) The genotype of the female offspring was $Foxp3^{WT/\Delta 2}$ and fed by C57BL/6J or Balb/c mother. The expression of RORγt in Tregs cells of colonic lamina propria of offspring mice fed by C57BL/6J strain and Balb/c strain were analyzed by flow cytometry. (**E**) Proportion of RORγt$^{+}$ Tregs cells in WT Tregs cells and Foxp3Δ2 Tregs cells of the two group.

The data represent the mean ± SD. The results are representative of at least three independent experiments with n ≥ 3. ns, no significance, *P < 0.05, **P < 0.01, ***P < 0.001, paired t test.

**Fig. S6**

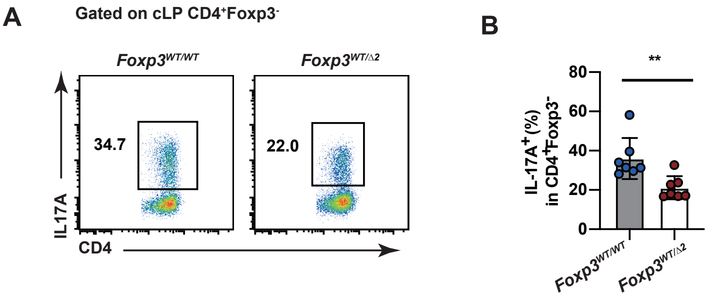
A
Gated on cLP CD4+Foxp3-
Foxp3WT/WT
Foxp3WT/Δ2
34.7
22.0
IL17A
CD4
B
**
IL-17A+(%) in CD4+Foxp3-
80
60
40
20
0
Foxp3WT/WT
Foxp3WT/Δ2

**Figure S6. Decreased Th17 response in cLP of *Foxp3*$^{WT/\Delta2}$ mice**

(**A**) After 14 days of *Citrobacter rodentium* infection, the expression of IL-17A in the cLP was detected by flow cytometry. (**B**) The proportion of Th17 cells in $CD4^{+}$ T cells.

The data represent the mean ± SD. The results are representative of at least three independent experiments with $n \geq 3$. ns, no significance, $^{*}P < 0.05$, $^{**}P < 0.01$, $^{***}P < 0.001$, unpaired t test.
